# Addition of chemotherapy to radiotherapy promotes progenitor-exhausted CD8⁺ T-cell clonal dominance in head and neck cancer

**DOI:** 10.64898/2026.03.10.710795

**Authors:** Charleen Chan Wah Hak, Anton Patrikeev, Antonio Rullan, Emmanuel C Patin, Victoria Roulstone, Lisa C Hubbard, Matthew Guelbert, Elizabeth S Appleton, Shane Foo, Isaac Dean, Amy Burley, Joan N Kyula-Currie, Holly Baldock, Jen Y Lee, Pablo Nenclares, Hannah Nanapragasam, Carmen Murano, Malin Pedersen, Maggie Chon U Cheang, Shree Bhide, Masahiro Ono, Kevin J Harrington, Alan A Melcher

## Abstract

Concomitant chemoradiotherapy (CRT) is a standard-of-care for unresectable locally-advanced head and neck squamous cell carcinoma (LA-SCCHN), but its immune effects, particularly compared to radiotherapy (RT) alone, remain unclear. Using syngeneic murine models, we integrated Nr4a3-Tocky reporter analysis with single-cell transcriptomics and T-cell receptor clonotyping comprehensively to profile intratumoural CD8⁺ T-cells following RT/CRT. We show that CRT uniquely drives robust antigen-specific clonal expansion and biases differentiation toward progenitor (or precursor) exhausted (T_PEX_) phenotypes, while RT favours terminal exhaustion (T_EX_). Single-cell analyses reveal CRT-induced clonal dominance within T_PEX_ subsets, suggesting the potential for enhanced immune reinvigoration. In peripheral blood mononuclear cells (PBMCs) from patients treated with CRT, high levels of T_EX_ cells were found early and at 3 months post-treatment, with delayed peripheral T_PEX_ expansion at 3 months, indicating phased progenitor recovery. These findings demonstrate distinct immunological remodelling by CRT versus RT and underscore the critical importance of treatment timing for optimising combination immunotherapy strategies in LA-SCCHN.

## INTRODUCTION

The interplay between tumour immunity and standard-of-care cancer therapeutics has become a major theme of oncological research. Central to this is the concept of the cancer-immunity cycle, a dynamic and potentially self-amplifying process through which tumour cell death fuels antigen release, dendritic cell (DC) activation, and subsequent T-cell priming, culminating in iterative cytotoxic responses against malignant cells (1). Yet, within the tumour microenvironment (TME), chronic antigen stimulation and immunosuppressive signals often derail T-cell differentiation, accelerating the emergence of dysfunctional, exhausted CD8⁺ T-cell phenotypes (2). T-cell exhaustion, classically defined by impaired effector function and upregulation of inhibitory receptors such as PD-1, LAG-3, and TIM-3, represents a major barrier to durable tumour control (3). This state is generally seen as irreversible, particularly in the terminally exhausted (T_EX_) subset. However, the less differentiated precursor exhausted (T_PEX_) population, marked by the transcription factor TCF1, retains proliferative potential and responsiveness to immune checkpoint inhibitors (ICIs), positioning it as a key target for therapeutic approaches that seek to reinvigorate anti-tumour immunity (4–6).

While the spotlight in recent years has deservedly fallen on ICIs, an important paradigm shift has emerged: radiotherapy (RT) and chemoradiotherapy (CRT) – historically viewed as being immunosuppressive – can also profoundly modulate the immune landscape in an immunostimulatory fashion. Indeed, clinical successes, such as the PACIFIC trial, illustrate the potential of RT and CRT to synergise with immunotherapy (7). RT and CRT can trigger immunogenic cell death (ICD), enhancing tumour antigen availability and inflammatory cytokine release (8). This process releases damage-associated molecular patterns (e.g., ATP, HMGB1, calreticulin), which activate both surface and intracellular pattern recognition receptors (PRRs) expressed by dendritic cells (DCs) and other antigen-presenting cells. These effects may transform the irradiated tumour into an *in-situ* vaccine capable of priming systemic, tumour-specific immune responses. Furthermore, recent studies have shown that CRT reshapes T-cell receptor (TCR) clonality by selectively expanding dominant T-cell clones associated with tumour specificity in head and neck, cervical, and rectal carcinomas (9–11). However, the functional consequences of this expansion remain unclear.

The incidence of squamous cell carcinoma of the head and neck (SCCHN) is rapidly rising in the developed world and it ranks as the seventh commonest cancer diagnosis worldwide (12). Definitive CRT remains standard-of-care for patients with unresected locally-advanced (LA) disease. Although the addition of cisplatin increases toxicity compared to RT alone, it improves local control and overall survival (13). The benefit of platin-based CRT is statistically significant and clinically meaningful, but relatively modest (approximately 6.5%) and only clearly apparent with meta-analysis of data from >100 trials (13). Attempts at de-escalation, through replacement of cisplatin with cetuximab, in HPV-associated oropharyngeal cancer have failed to reduce toxicity and have, instead, shown worse survival outcomes (14, 15), reinforcing the primacy of CRT as standard-of-care for unresected LA-SCCHN. Nonetheless, de-escalation strategies are still being investigated in select HPV-positive oropharyngeal cancer cohorts. For example, the ECOG-ACRIN 3311 trial showed that, in surgically resected intermediate-risk patients, adjuvant RT could be safely reduced from 60 Gy to 50 Gy without compromising disease control (16). In parallel, ^18^F-fluoromisonidazole (FMISO) PET has been shown to identify hypoxia-negative tumours suitable for RT dose reduction, achieving excellent outcomes even with de-escalated CRT doses (∼30 Gy), supporting its use as a potential functional biomarker to guide adaptive RT dose and volume reduction (17). Despite these advances in refining locoregional therapy, overall survival outcomes for patients with LA-SCCHN have plateaued, highlighting the need for novel strategies that move beyond optimisation of CRT parameters alone.

One promising avenue has been the integration of ICI with standard treatment backbones. However, the failure to improve outcomes by combining ICI directly with (C)RT, as evidenced in multiple phase III trials, such as JAVELIN Head and Neck 100 (18), KEYNOTE-412 (19), GORTEC 2017-01 (20), IMvoke010 (21) and COMPARE (22), underscores a critical gap in our understanding of how standard therapies shape the immune landscape. Preclinical models have almost exclusively focused on combining ICI with RT alone, failing to characterise potential differences in immunologic effects of CRT. While CRT is widely used and clinically effective, its immunomodulatory impact relative to RT, including how it influences T-cell differentiation trajectories, precursor exhaustion, and clonal dynamics, has largely been ignored. Notably, antigen persistence during CRT could promote both expansion and premature exhaustion of TILs, particularly in highly inflammatory environments like SCCHN (23). Emerging evidence underscores the prognostic and functional relevance of different CD8⁺ T-cell exhaustion states in SCCHN. Higher frequencies of polyfunctional CD8+ T_EX_ were associated with improved overall and recurrence-free survival in one study (24), whereas another showed that CD8^+^ T_PEX_ subset enrichment correlated with favourable outcomes (23). Transcriptional analyses implicate TCR signalling as a key determinant of exhausted T-cell responsiveness to neoadjuvant immunotherapy in SCCHN (25).

Therefore, we used syngeneic immunocompetent animal models to show that while both RT and CRT increase CD8⁺ T-cell infiltration, only CRT depends on these cells for therapeutic efficacy. We show that CRT drives more focused clonal expansion and effector differentiation of antigen-engaged CD8⁺ T-cells, with a preferential shift toward effector memory and T_PEX_, rather than T_EX_, states. Translational analyses of patient PBMCs revealed systemic immune modulation, with CRT associated with stronger T_EX_ accumulation and a delayed rise in putative T_PEX_ subsets compared with RT. Together, these findings define distinct immunologic consequences of RT and CRT, inform biomarker development, and provide a roadmap for rational integration of immunotherapy.

## RESULTS

### Radiotherapy (RT) and chemoradiotherapy (CRT) both increase CD8^+^ T-cell infiltration into tumours, but the therapeutic effect is CD8^+^ T-cell dependent only with CRT

Compared to RT, CRT yielded a modest, non-significant improvement in tumour control in both mEER and MOC1 preclinical models (Fig. 1A, S1A, S2A), in line with the relatively modest benefits seen with meta-analysis in clinical studies (13). Survival was significantly prolonged in the mEER model (Fig. 1B), but not in MOC1 tumours (Fig. S2B). In the mEER model, CD8^+^ tumour-infiltrating lymphocyte (TIL) dynamics varied over time following RT as determined by flow cytometry from baseline: numbers declined at day 3 post-RT, consistent with radiation-induced intratumoural lymphopenia, recovered by day 7 post-RT, and peaked at day 10 (Fig. 1C). Both RT and CRT increased CD8^+^ TIL abundance compared to vehicle or cisplatin-alone controls; however, no significant differences were observed between RT and CRT at day 10 or day 14 following treatment when quantified by immunohistochemistry (Fig. 1D) and flow cytometry (Fig. 1E). Furthermore, there were no significant differences in the percentage of PD-1^hi^ expressing CD8^+^ T-cells between RT and CRT treatment groups (Fig. 1F).

**Fig. 1:**
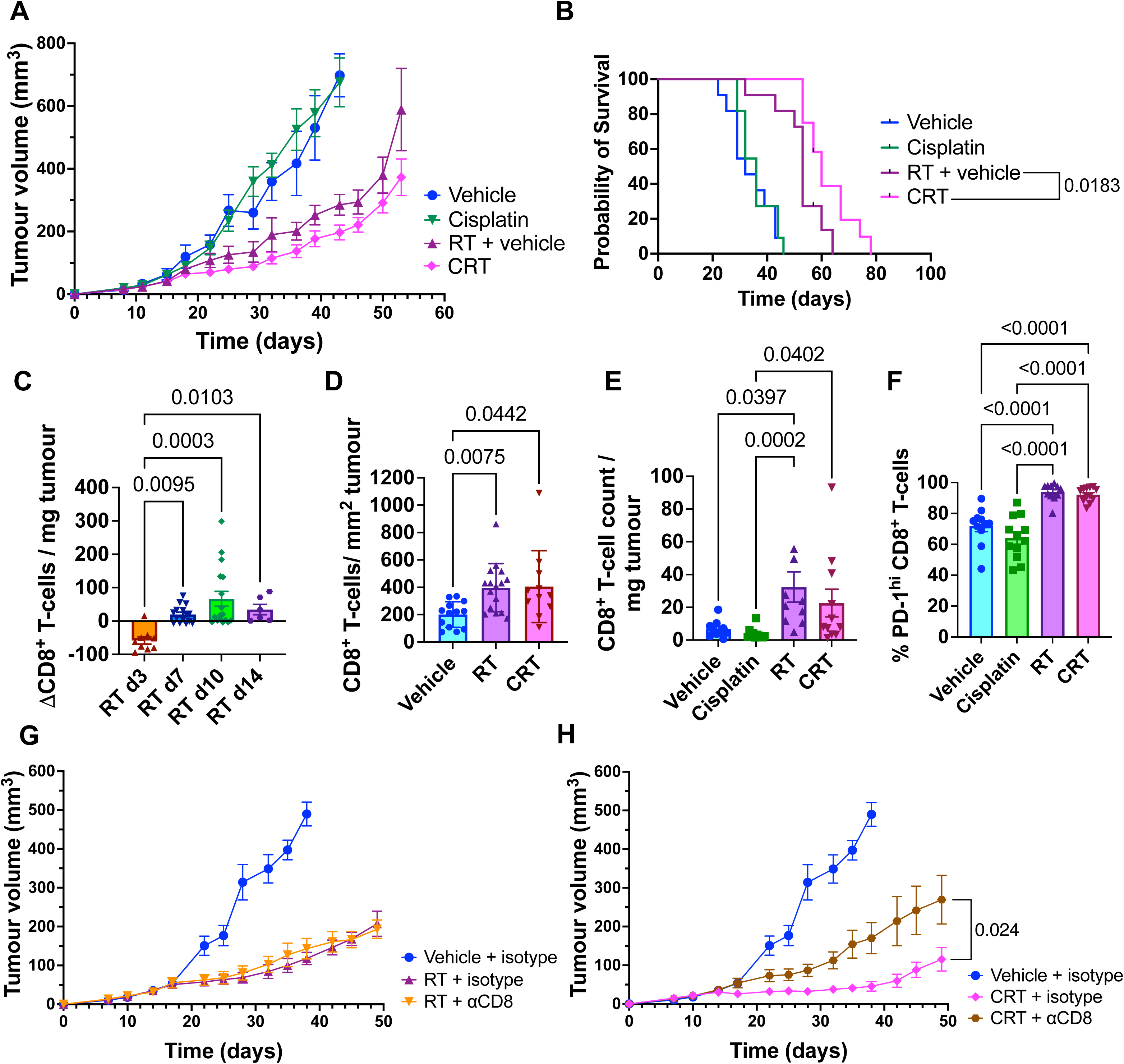
Both radiotherapy (RT) and chemoradiotherapy (CRT) increase CD8^+^ T-cell infiltration into tumours, but therapy is only CD8^+^ T-cell dependent in CRT. Experiments performed in the mEER model. A. Tumour growth curves across indicated treatment conditions. B. Survival curves across indicated treatment conditions (A-B: Vehicle n=11, Cisplatin n=11, RT + vehicle n=11, CRT n=12; 2 independent experiments). C. Absolute change of CD8^+^ T-cell counts/mg tumour vs average of timematched controls across indicated timepoints post-RT as measured by flow cytometry (RT d3 n = 9, RT d7 n = 15, RT d10 n = 17, RT d14 n = 6; 6 independent experiments of 2 consecutive timepoints). D. CD8^+^ T-cells/ mm^2^ tumour at 14 days post-RT by immunohistochemistry, necrotic regions excluded, quantified by QuPath v. 0.3.0. (Vehicle n=13, RT, n=16, CRT n=11; 3 independent experiments). E. CD8^+^ T-cell count/ mg of tumour by flow cytometry (Vehicle n=10, Cisplatin n=11, RT n=10, CRT n=11; 2 independent experiments). F. Percentage of PD-1^hi^ CD8^+^ T-cells by flow cytometry (Vehicle n=11, Cisplatin n=12, RT n=11, CRT n=12; 2 independent experiments). G-H: Tumour growth curves of ⍺CD8 depletion experiments, with mice receiving either RT (G) or CRT (H) (Vehicle + isotype n=6, RT + isotype n=6, RT + ⍺CD8 n = 5, CRT + isotype n = 6, CRT + ⍺CD8 n = 6; data from 1 experiment). Results are means ± SEM and *n* denotes mice per group. Parametric statistics were only applied to normally distributed data (Shapiro-Wilk test). P-values shown were determined by the log-rank (Mantel–Cox) (B), Kruskal Wallis test with Dunn’s multiple comparisons test (C-E), Ordinary one-way ANOVA with Tukey’s multiple comparisons test (F) and Two-tailed Unpaired t test (H).

To assess the function of CD8^+^ T-cells, we performed depletion studies under RT and CRT conditions. In the mEER model, RT-mediated tumour control was CD8^+^ T-cell-independent (Fig. 1G, Fig. S1B), whereas CRT (cisplatin + RT) showed CD8^+^ T-cell-dependence, as depletion significantly decreased tumour control (Fig. 1H, Fig. S1B, Fig. S3). These findings indicate a mechanistic role for CD8^+^ T-cells in CRT response in this model. In the MOC1 model, which exhibits higher baseline CD8^+^ T-cell infiltration, depletion affected neither RT nor CRT (Fig. S2D-E). Subsequent experiments focused on the mEER model, where CRT conferred a modest survival benefit [in line with the clinical meta-analysis (13)], to investigate how addition of cisplatin to RT modulates CD8^+^ T-cell function in this setting.

### CRT reduces the proportion of persistently engaged CD8^+^ TILs compared to RT

Immunoprofiling via flow cytometry of mEER tumours showed that RT and CRT broadly enhanced lymphocyte activation, proliferation, and cytotoxicity, with CRT exerting the stronger effect overall (Fig. S4). Both treatments increased CD44 expression (Fig. S4A) and expanded granzyme B^+^/perforin^+^ cytotoxic populations (Fig. S4B) across CD8⁺ T-cell, NK, and NKT subsets. CRT significantly increased Ki67^+^ proliferating CD8⁺ T and NK cells (Fig. S4C), as well as T-bet^+^ (Fig. S4D) expression in Tconv and CD8⁺ T-cells. Effector memory (Fig. S4E) and PD-1⁺/TIM-3⁺ checkpoint-positive (Fig. S4F) CD8⁺ T-cells were also increased by both treatments.

As cisplatin-based CRT was CD8^+^ T-cell-dependent, whereas RT alone was not, we next asked whether CRT differentially affects the quality or kinetics of T-cell antigen engagement (Fig. 2A). To address this, we employed the Nr4a3-Tocky (“Timer of cell kinetics and activity” reporter system) (26–28). This system utilises a Fluorescent Timer (FT) protein that undergoes a spontaneous, time-dependent shift in Timer fluorescence following TCR-mediated induction of Nr4a3, a TCR downstream nuclear receptor gene. The FT protein is initially expressed as an immature blue fluorophore, which matures into a stable red fluorophore with a half-life of approximately 4 hours (27). Consequently, newly antigen engaged T-cells express blue fluorescence (“new”), persistently engaged T-cells express both blue and red (“persistent”), and cells in which TCR signalling has ceased (“arrested”) express red fluorescence only.

**Fig. 2:**
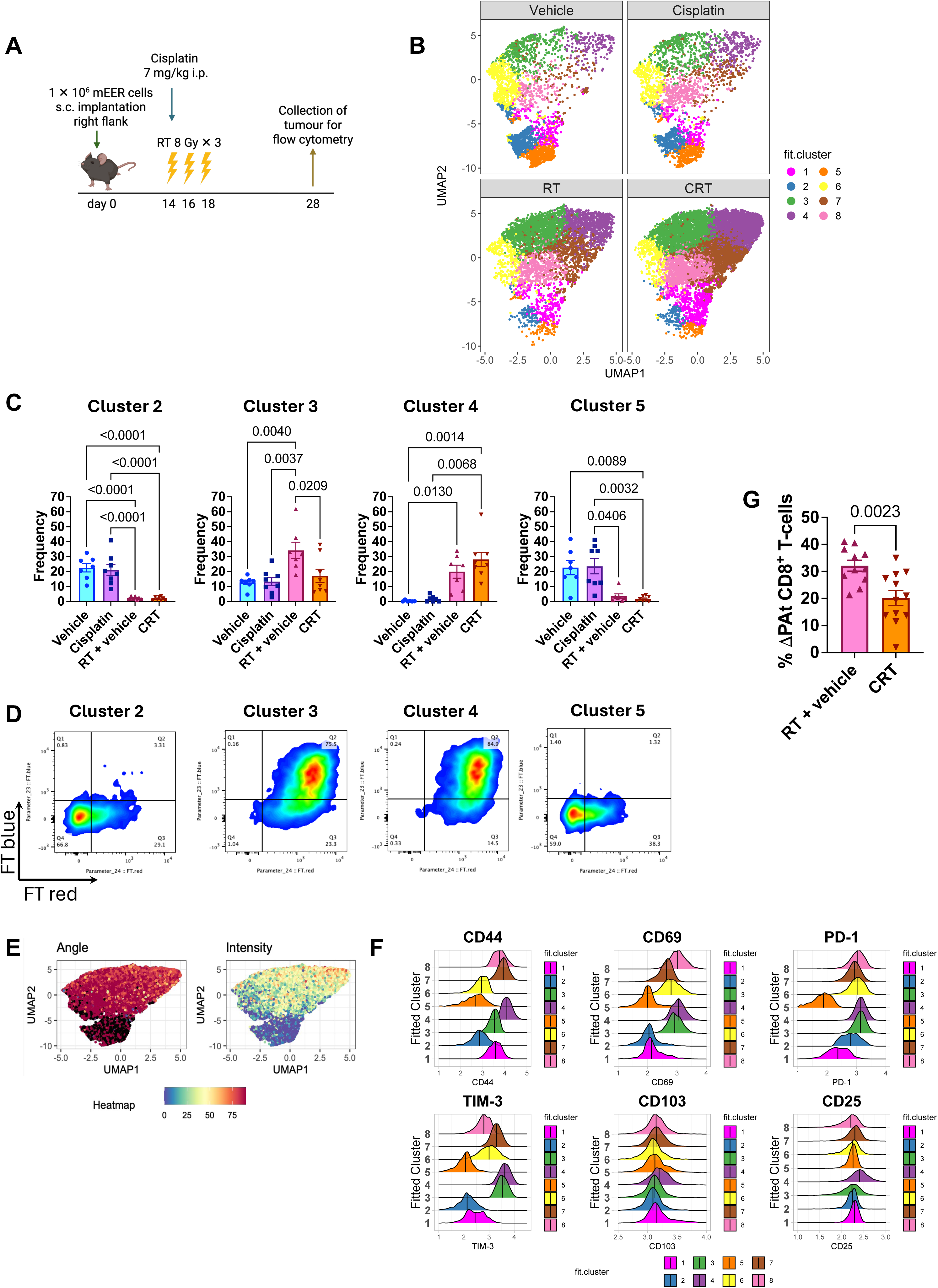
T-cell dynamics reveals differences in tumour T-cell populations between RT and CRT, with a reduced percentage increase in persistently engaged CD8^+^ T-cells in CRT versus RT. **A.** Schematic of the treatment timeline in Nr4a3-Tocky mEER tumour-bearing mice. *Created with Biorender.com.* **B-F:** UMAP analysis of CD8^+^ tumour-infiltrating lymphocytes (TILs), results concatenated from 30 mice (Vehicle n=7, Cisplatin n=8, RT + vehicle n=7, CRT n=8; 1 experiment). **(B)** UMAP plot showing distribution of clusters 1–8 across treatment conditions. **(C)** Selected clusters 2, 3, 4 and 5 showing significant differences between treatment groups (Vehicle n=7, Cisplatin n=8, RT + vehicle n=7, CRT n=8). **(D)** Timer fluorescence patterns within clusters 2, 3, 4 and 5. Cluster 3 is defined by a persistent-arrested (PAt) Tocky profile with high activation and immune checkpoint expression. **(E)** Heatmap overlays of Tocky “angle” (antigen engagement), “intensity” (TCR signalling strength). **(F)** Histograms illustrating marker expression by cluster. **G.** Percentage change in PAt CD8^+^ T-cells (% ΔPAt) in RT + vehicle and CRT groups versus the average of vehicle group (RT + vehicle n=11, CRT n=12; 2 independent experiments). Results are means ± SEM and *n* denotes mice per group. Parametric statistics were only applied to normally distributed data (Shapiro-Wilk test). P-values shown were determined by Ordinary one-way ANOVA with Tukey’s multiple comparisons test (E: cluster 2-3), Kruskal Wallis test with Dunn’s multiple comparisons test (E: cluster 4-5) and Two-tailed Unpaired t test (G). Significant outliers were removed via ROUT test (Q = 1%). All data are representative of two independent experiments.

These dynamics are quantified using the ‘Timer Angle’, a trigonometric transformation where 0° (blue only) represents the onset of activation, 45° (blue and red) indicates sustained persistent signaling, and 90° (red only) identifies the cessation of TCR engagement. Additionally, the overall fluorescence intensity provides a qualitative measure of cumulative TCR signal strength.

To evaluate treatment-induced remodeling of the immune landscape, UMAP analysis was performed on concatenated CD8⁺ TIL data from all treatment groups (Fig. 2B). Both treatments increased the abundance of antigen-engaged, activated (CD44^hi^ CD69^hi^ CD25^hi^) and immune checkpoint-expressing (PD-1^hi^ TIM-3^hi^) CD8^+^ TILs compared to vehicle or cisplatin alone (Fig. 2C-F). This change was reflected in a significant increase in the frequency of cluster 4 (Fig. 2C), which was predominantly composed of cells exhibiting high Timer Intensity and ‘persistent’ TCR signal (Fig. 2D). Conversely, there was a corresponding decrease in the frequency of clusters 2 and 5 (Fig. 2C), which were dominated by a profile of timer-negative status, low activation and low immune checkpoint expression (Fig. 2D-F).

Focusing on the comparison between RT and CRT, a notable difference emerged within cluster 3. CRT-treated tumours exhibited a significant decrease in this cluster (Fig. 2C), which was defined by a “persistent-arrested” antigen engagement profile – characterised by sustained TCR signalling – despite displaying activation and immune checkpoint marker expression comparable to cluster 4 (Fig. 2D-F). Overall, CRT did not enhance overall antigen engagement compared to RT but, instead, specifically reduced the proportion of “persistent-arrested” (PAt) CD8^+^ TILs relative to vehicle controls (Fig. 2G). This subset may represent terminally exhausted cells, secondary to continuous antigen engagement (3, 29, 30).

Interestingly, addition of anti-PD-1 antibody (⍺PD-1) had no effect with RT alone (Fig. S5A), while there was a modest, but statistically non-significant, improvement in tumour control with ⍺PD-1 in CRT-treated mice (Fig. S5B). These findings suggest differential impacts of RT and CRT on T cell exhaustion, motivating further interrogation of the transcriptional and phenotypic states of antigen-engaged TILs using single-cell approaches.

### CRT increases antigen-engaged TILs with effector memory (T_EM_) relative to naïve phenotype (T_N_), when compared to RT alone

To investigate the impact of CRT on T-cell phenotypes in more detail, we combined the Nr4a3-Tocky system with single-cell transcriptomic and protein profiling, using AbSeq, and TCR sequencing (Fig. 3A). Tumours and tumour-draining lymph nodes (TDLNs) were harvested from mice treated with vehicle, RT, or CRT, and cells were analysed by UMAP clustering based on AbSeq expression profiles (Table S3). Within the tumour, TILs were dominated by effector memory (T_EM_) phenotype (CD62L^-^ CD44^+^), whereas T-cells from TDLNs were primarily naïve (T_N_; CD62L^+^) and central memory (T_CM_; CD62L^+^, CD44^+^) (Fig. 3B-E).

**Fig. 3:**
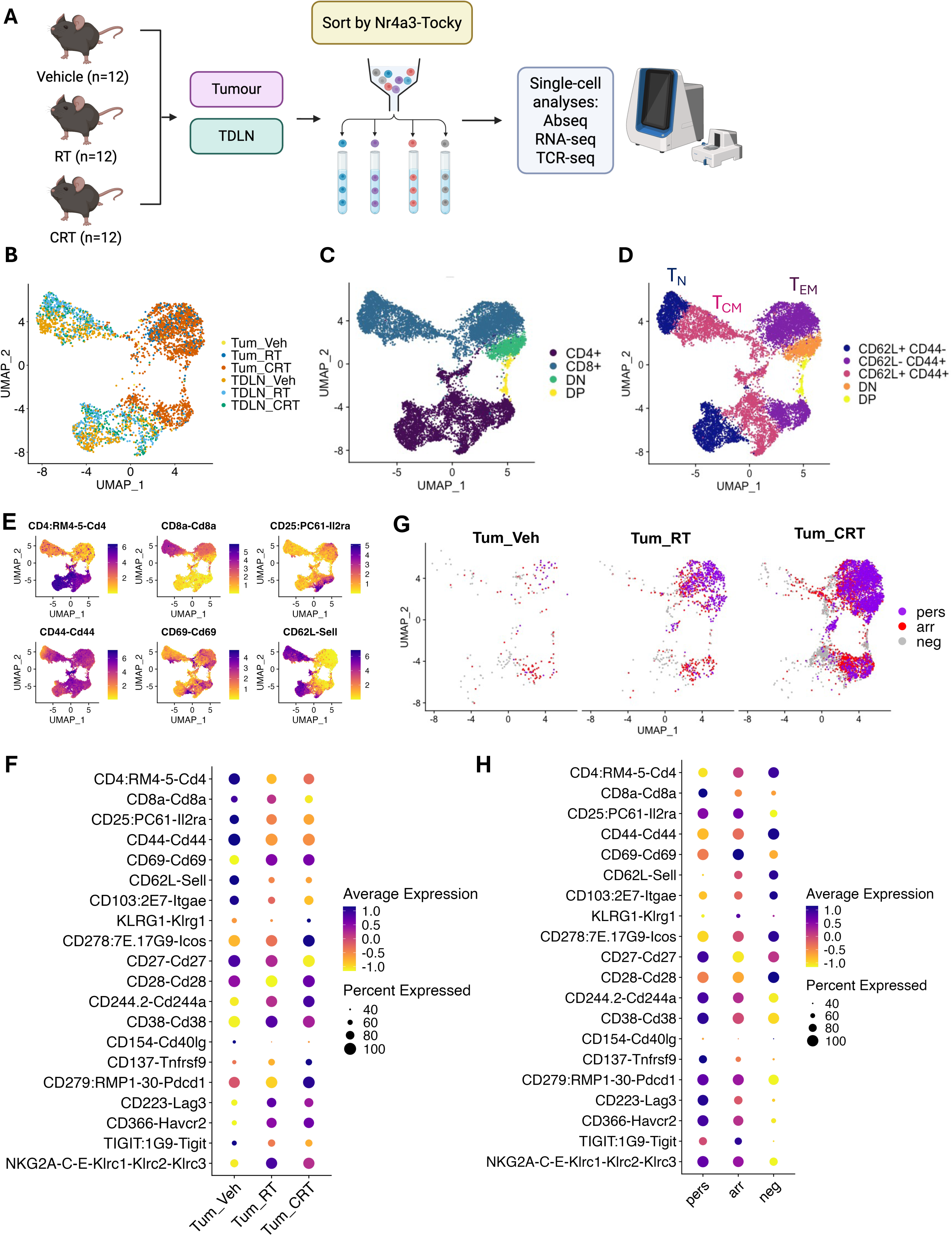
CRT increases antigen-engaged TILs with effector memory over naïve phenotype compared to RT; persistently engaged T-cells show higher inhibitory checkpoint expression linked to exhaustion. **A.** Experimental schema: Single-cell analysis was performed on tumour-infiltrating lymphocytes (TILs) from tumours (Tum) or TDLN harvested from mEER-bearing Nr4a3-Tocky mice treated with vehicle (Veh), RT or CRT. Cells were sorted according to Tocky antigen-engagement profile prior to integrated single-cell analyses (Vehicle n=12, RT n=12, CRT n=12; 1 experiment). **B-H:** Cells were subject to UMAP clustering based on Abseq expression profile. **(B)** UMAP by sample tag. **(C)** Cell annotation of T-cell subtype: CD4, CD8, DN (double negative) and DP (double positive). **(D)** Functional T-cell subtype annotation based on expression markers: T_N_ (naïve, CD62L^+^), T_CM_ (central memory, CD62L^+^, CD44^+^), T_EM_ (effector memory, CD62L^-^ CD44^+^). **(E)** Heatmap of selected Abseq marker expression overlaid on UMAP. **(F)** Dot plot of Abseq marker expression in TILs by treatment. **(G)** Distribution of Tocky populations: “persistent” (pers), “arrested” (arr) and “timer negative” (neg) within the UMAP space. **(H)** Dot plots of Abseq marker expression across different Tocky populations in TILs.

Within TILs, following both RT and CRT, there was increased expression of activation markers (CD69, CD38), co-stimulatory receptors (CD278 [ICOS], CD137 [4-1BB]) and markers of inhibition/exhaustion (CD223 [LAG3], CD366 [TIM-3]), TIGIT and NKG2A-C-E) compared to vehicle. There was also a reduction in the average expression of the activation/Treg marker, CD25 compared to vehicle (Fig. 3F). Again, focusing particularly on differences with the addition of chemotherapy to RT, compared to RT alone, CRT showed increased KLRG1, CD278 (ICOS), CD28 and CD244a expression and reduced CD27 and CD38 expression. This profile is consistent with chemotherapy enhancing the induction of T_EM_ relative to naïve T-cells, compared to RT alone.

### Persistently-engaged T-cells express higher levels of multiple inhibitory checkpoint molecules consistent with continual antigen engagement driving T-cells towards exhaustion

To assess the temporal dynamics of antigen engagement across treatment groups, we mapped Tocky-defined populations (“persistent” [pers], “arrested” [arr], and “timer-negative” [neg]) onto the UMAP landscape (Fig. 3G). Untreated tumours (Tum_Veh) exhibited sparse antigen-engaged T cells, with a predominance of timer-negative cells. In contrast, both RT and CRT markedly induced a striking predominance of antigen-engaged TILs within the TME. This expansion was most pronounced following CRT, which demonstrated a dense and cohesive accumulation of “persistent” (pers) T-cells (purple in Fig. 3G), indicative of sustained TCR signalling.

Examination of marker expression across Tocky populations showed that “persistent” T-cells expressed markedly higher levels of multiple immune checkpoint markers: CD279 (PD-1), CD233 (LAG-3), CD366 (TIM-3) and TIGIT, and the exhaustion-associated markers, CD244a and NKG2A-C-E, compared with other Tocky loci (Fig. 3H). This profile is consistent with continual antigen engagement driving T-cells towards an exhausted phenotype (3). In contrast, “timer-negative” T-cells demonstrated relatively higher expression of CD62L, CD103, CD27 and CD28, indicative of a less antigen-experienced or memory-associated state (Fig. 3H). Interestingly, while the “arrested” Tocky population displayed intermediate levels of immune checkpoints (e.g. PD-1, LAG-3, TIM-3), they reached a positive peak in CD69 expression and a negative peak in CD27. Furthermore, the inclusion of KLRG1-high cells within the arrested population suggests that these cells sustain enhanced cytotoxic potential (Fig. 3H). These dynamics suggest that T-cells with historical antigen engagement undergo an active remodelling phase at the Arrested stage, leaving the terminal exhaustion state observed in persistently stimulated cells.

While the single-cell transcriptomic analysis demonstrates increased overall antigen engagement following CRT, the persistent-arrested (PAt) subset defined by Nr4a3-Tocky dynamics cannot be directly identified within the scRNA-seq dataset (Fig. 3G). However, flow cytometry using the Tocky reporter revealed that CRT reduced the proportion of PAt cells compared with RT alone (Fig. 2G). Together, these findings indicate that CRT is associated with increased overall T-cell antigen engagement while reducing the accumulation of more exhausted PAt cells, relative to RT alone.

### CRT leads to more focused, clonally expanded CD8^+^ T-cell populations with an effector profile

Clonal expansion occurred almost exclusively within tumours treated with RT and CRT, with CRT leading to greater clonal expansion than RT alone (Fig. 4A-D). Interestingly, hyperexpanded clones were made up of only three paired TCR α/β CDR3 sequences, that occurred exclusively in T-cells from tumours treated with CRT, highlighting striking clonal dominance (Fig. 4B). Following downsampling to the minimum clonotype count for comparability, consistently lower diversity in TILs from CRT-versus RT-treated tumours was still found by various indices (Fig 4E). For reference, Shannon and Simpson indices combine clonotype richness and evenness, Chao1 and ACE indices estimate unseen diversity and the Inverse Pielou index was used to assess clonal skewing (31). Hence, the TCR repertoire in the CRT group was less even than RT and more clonal/skewed towards dominant clones. With increasing clonal expansion, a progressive predominance of CD8⁺ over CD4⁺ T-cells was observed, with hyperexpansion occurring exclusively within the CD8⁺ compartment (Fig. 4F).

**Fig. 4:**
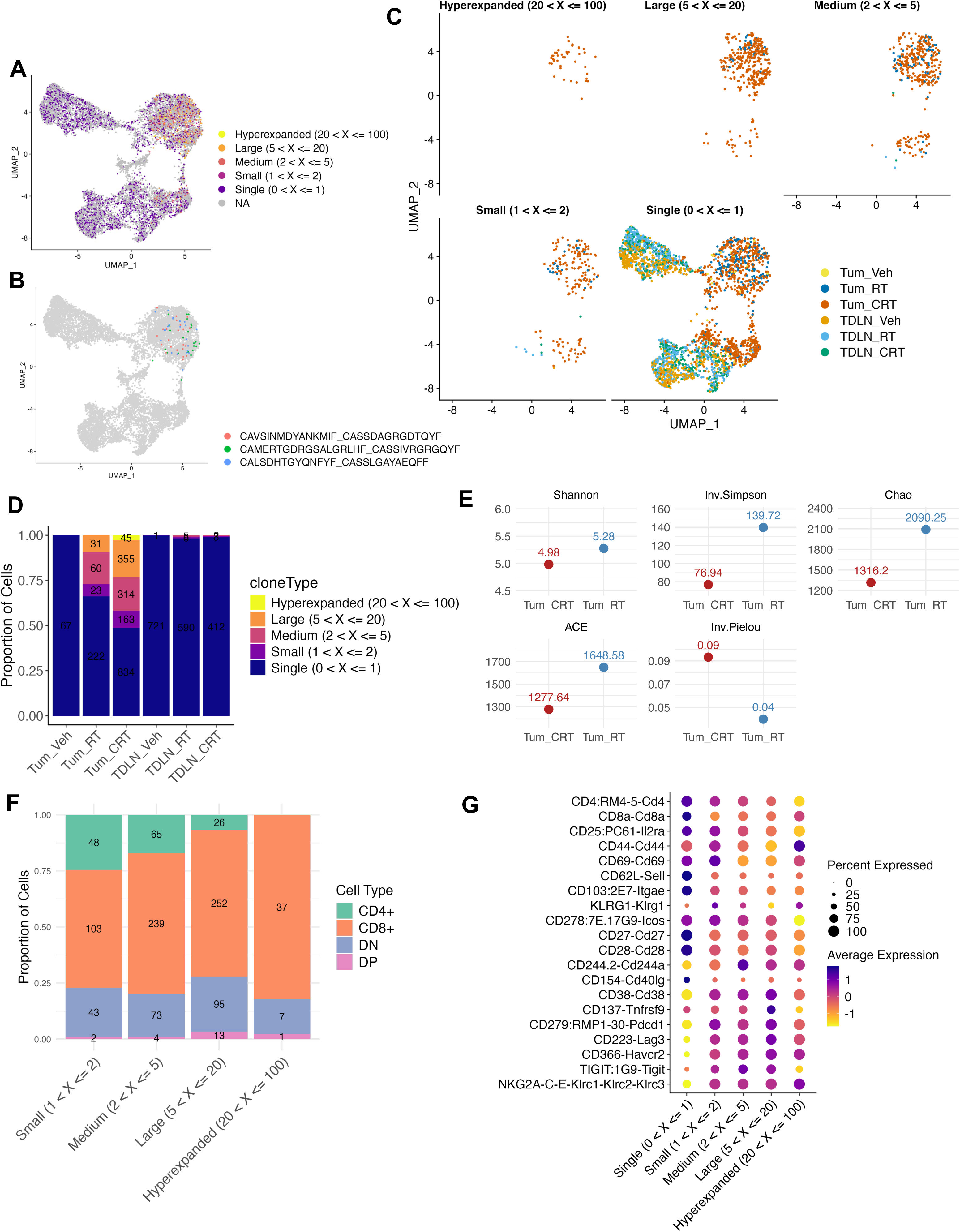
CRT leads to more focused, expanded CD8+ T-cell populations with an effector profile than RT. Single-cell TCR sequencing was performed on sorted tumour-infiltrating lymphocytes (TILs) from tumours (Tum) or TDLN harvested from mEER-bearing mice treated with vehicle (Veh), RT or CRT (Vehicle n=12, RT n=12, CRT n=12, 1 experiment). **A.** Distribution of T-cell clonal expansion within UMAP space, categorised by clone frequency within the total TCR repertoire: hyperexpanded (>20 and ≤100), large (>5 and ≤20), medium (>2 and ≤5), small (>1 and ≤2) and single (>0 and ≤1). Cells without information on paired TCR α (alpha) and β (beta) chain CDR3 (Complementarity-Determining Region 3) sequence information are denoted by “NA”. **B.** Three CDR3 sequences which were hyperexpanded. **C.** Distribution of categories of clonal expansion within UMAP space by sample type and treatment group. **D.** Categories of clonal expansion by sample type and treatment group by proportion of cells **E.** Clonal diversity dotplots comparing various diversity indices between Tum_RT and Tum_CRT: Shannon; Inverse (Inv) Simpson; Chao1 (Chao) Abundance-based Coverage Estimator (ACE) and Inv Pielou. **F.** Proportions of T-cell type (CD4+, CD8+, double negative [DN], double positive [DP]) by categories of clonal expansion. **G.** Abseq expression marker profile across different clonal expansion categories.

Hence, CRT reshaped the TCR repertoire of TILs and altered the expression profile of hyperexpanded CD8^+^ T-cells. In all singleton clones across treatment conditions, there was a predominance of naïve or early memory (CD44^lo^, CD62L^hi^, CD27^hi^, CD28^hi^) and/or tissue-resident T-cells (CD103^hi^) (Fig. 4G). When the expression profile of the hyperexpanded clones (seen only with CRT) was examined, relatively higher expression of the activation marker CD44 was observed, with relatively lower multiple immune-checkpoint /inhibitory markers – CD279 (PD-1), CD223 (LAG-3), TIGIT and CD38, compared to the other clonal subgroups (Fig. 4G). These findings suggest that CRT-induced hyperexpanded CD8^+^ T-cells predominately acquire an effector-like, rather than exhausted, phenotype.

### CRT leads to greater clonal expansion of antigen-engaged CD8^+^ T-cells than RT

Correlation of TCR-seq data with the Tocky profile revealed that CRT induced greater clonal expansion of antigen-engaged tumour-infiltrating CD8⁺ T cells (“persistent” and “arrested”) compared to RT, whereas vehicle-treated tumours showed minimal clonal expansion (Fig. 5A–B). In contrast, TILs lacking evidence of antigen engagement at the time of analysis (“timer-negative”) demonstrated limited clonal expansion across treatment groups.

**Fig. 5:**
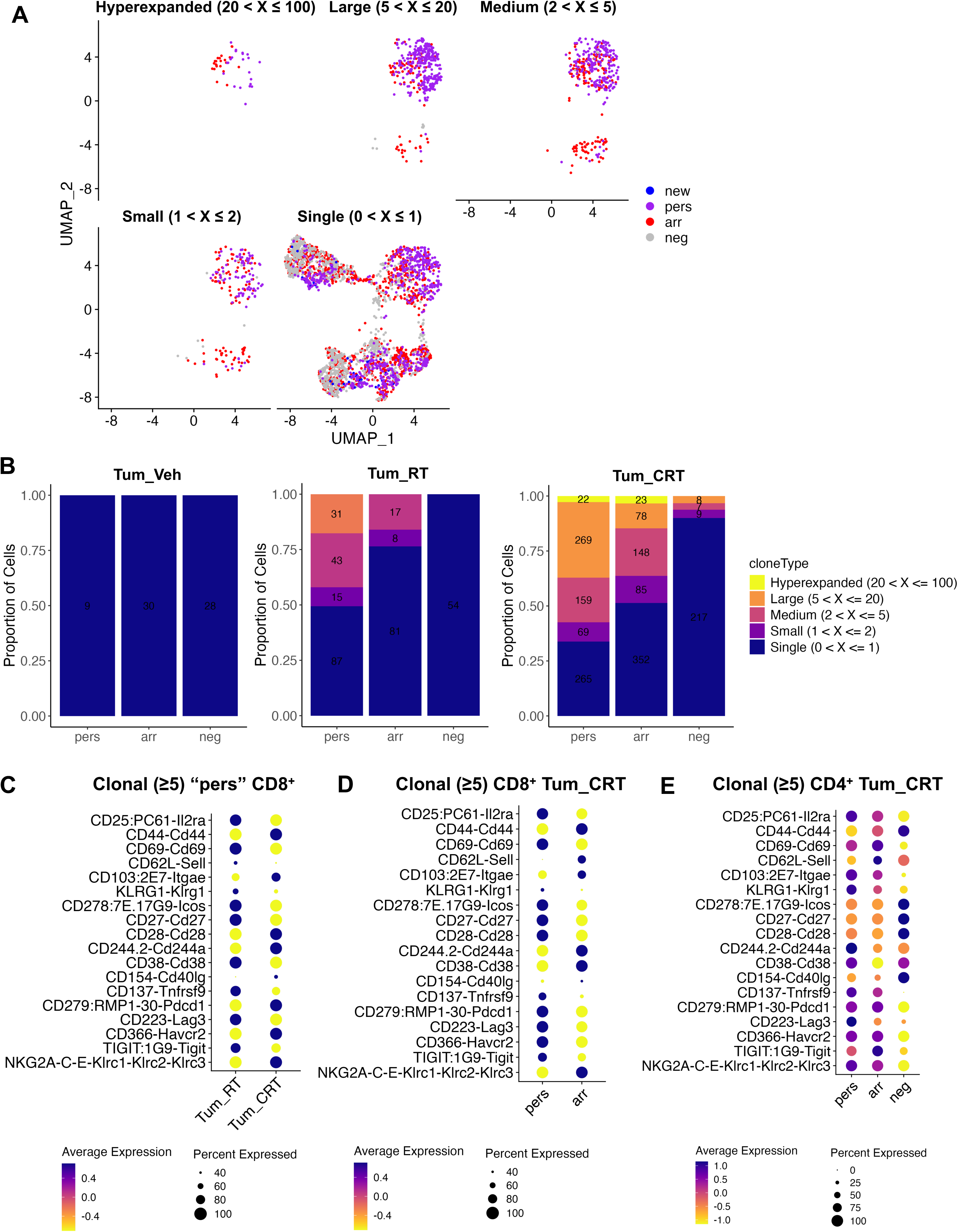
CRT leads to greater clonal expansion of antigen-engaged T-cells than RT, and clonal CD8 TILs treated with CRT show relative acquisition of a less exhausted phenotype on transition from persistent (“pers”), to historical, T-cell engagement (“arr”). Single-cell analysis was performed on sorted tumour-infiltrating lymphocytes (TILs) from tumours (Tum) or TDLN harvested from mEER-bearing mice treated with vehicle (Veh), RT or CRT (Vehicle n=12, RT n=12, CRT n=12, 1 experiment). **A.** Distribution of Tocky populations within UMAP space according to categories of clonal expansion: “new”, “persistent” (pers), “arrested” (arr) and “timer negative” (neg). **B.** Proportions of cells undergoing clonal expansion by Tocky population. **C-E:** Abseq marker expression profile of various TILs subsets: **(C)** Clonal (≥ 5) and “persistent” CD8^+^ TILs by treatment type, **(D)** Clonal (≥ 5) CD8^+^ TILs from mice treated with CRT (Tum_CRT) by Tocky population and **(E)** Clonal (≥ 5) CD4^+^ TILs treated with CRT (Tum_CRT) by Tocky population.

Given that Timer fluorescence reflects real-time TCR signalling dynamics, the enrichment of expanded clonotypes within the “persistent” compartment in CRT-treated tumours (Fig. 5B) suggests that a discrete subset of clones is either undergoing sustained proliferative expansion driven by continuous antigen stimulation or, alternatively, that once expanded, these clones preferentially occupy and remain within the persistently engaged niche.

Despite treatment-matched sampling, we observed limited clonal overlap between tumour and TDLN compartments (Fig. S6). Together with the apparent confinement of expanded clonotypes to defined Tocky signalling stages (Fig. 5B, S7), this supports a model in which CRT promotes spatially restricted, antigen-dependent clonal expansion within the TME.

### Clonal CD8^+^ TILs treated with CRT show relative acquisition of a less exhausted phenotype as they progress from “persistent” to “arrested”

To capture broader antigen-driven activity, the expression profile of specific clonal populations (frequency ≥5) was studied in relation to their Tocky profile in different samples. Firstly, clonal “persistent” CD8^+^ TILs were compared between tumours treated with RT and CRT; CD8^+^ TILs from CRT-treated tumours showed increased expression of activation and co-stimulation markers (CD44, CD28 and CD154 [CD40L]), the residency marker CD103, and antigen-experienced/ exhaustion markers (CD244a, PD-1, CD366 [TIM-3], NKG2A-C-E), as compared to RT-treated tumours (Fig. 5C). Overall, this suggests that continuously antigen-engaged CD8^+^ TILs shift towards a more activated, tissue-resident, and partially exhausted phenotype following CRT, rather than RT.

Next, all clonal CD8^+^ TILs with a frequency ≥5 from CRT-treated tumours were examined by Tocky profile; these were “persistent” versus “arrested”, as there were no sufficiently expanded cells in the “new” or “timer negative” subsets (Fig. 5D). Here again, consistent with the model of chronic antigen stimulation driving exhaustion, persistently antigen-engaged CD8^+^ TILs exhibited reduced activation (CD44) and increased expression of multiple immune checkpoints, CD279 (PD-1), CD223 (LAG-3), CD366 (TIM-3) and TIGIT, compared to historically antigen-engaged, but now “arrested”, cells. Moreover, a small percentage of “persistent” cells showed elevated levels of the terminal differentiation marker KLRG1 compared to “arrested”. Collectively, these findings suggest that progression along the Tocky trajectory from “persistent” to “arrested”, rather than sustained antigen re-engagement to maintain the “persistent” state, may mitigate T-cell exhaustion. Of note, this analysis could not be performed for TILs from RT-treated tumours, as there were no “arrested” clonal (frequency ≥5) CD8^+^ TILs present in this subset, only “persistent” cells.

Clonal CD4^+^ TILs, with a frequency ≥5, were only present in tumours treated with CRT and not RT (Fig. 5E). This is consistent with a model in which CD4^+^ T-cells support enhanced clonal expansion of CD8⁺ TILs following CRT more effectively than after RT alone. Similarities were observed in the expression profiles of “persistent” and “arrested” clonal CD4^+^ and CD8^+^ TILs; for example, “persistent” CD4^+^ TILs also exhibited higher expression of inhibitory/exhaustion-associated markers such as LAG3 and KLRG1, as well as activation markers CD25 and CD137 (4-1BB), compared to their “arrested” counterparts (Fig 5D-E). These are consistent with “persistent” cells representing a more chronically stimulated and potentially exhausted state in both CD4^+^ and CD8^+^ TILs.

### CRT compared to RT increases the proportion and clonal expansion of precursor exhausted (T_PEX_), relative to terminally exhausted (T_EX_), CD8+ TILs in the arrested phase

To define treatment-induced changes in T-cell states, sorted single T-cells were subjected to UMAP clustering based on their global transcriptomic profiles. Due to inadequate cell numbers and integrity, scRNAseq data was only available for “arrested” (arr) and “timer negative” (neg) samples, which nevertheless reflect either end of the Timer expression trajectory. UMAP clustering initially identified 11 transcriptionally distinct T-cell clusters. Based on shared lineage markers, characteristic gene expression of T_PEX_ and T_EX_ states in the published literature (4, 6, 32), and proximity in low-dimensional space, closely related clusters were consolidated into 9 annotated T-cell subsets.

As shown in Fig. 6A-B, naïve CD4⁺ and CD8⁺ T-cells were identified by high expression of CD62L (*Sell*). Classical CD8⁺ T_PEX_ cells were characterized by elevated TCF1 (*Tcf7*), intermediate PD-1 (*Pdcd1*), and low TIM-3 (*Havcr2*) and CD62L (*Sell*), indicative of antigen experience with preserved progenitor features (4, 6, 32). CD8⁺ T_EX_ cells expressed low TCF1, but high TOX, PD-1, LAG-3, and TIM-3, with interestingly maintained Granzyme B (*Gzmb*) and perforin (*Prf1*) expression expression. CD8⁺ cytotoxic (“cyto”) cells exhibited high expression of granzymes (*Gzma, Gzmb*) and perforin (*Prf1*), not falling into the T_PEX_ or T_EX_ phenotype. Two distinct FOXP3-enriched Treg-like subsets were identified: Treg1, displaying an effector/suppressive phenotype (high PD-1, CTLA-4, ICOS, IL2RB and ID2), and Treg2, with a more naïve/central-memory profile (high CD62L and IL7R). Both states correspond to previously described functional Treg phenotypes documented in regulatory T-cell heterogeneity studies (33, 34).

**Fig. 6:**
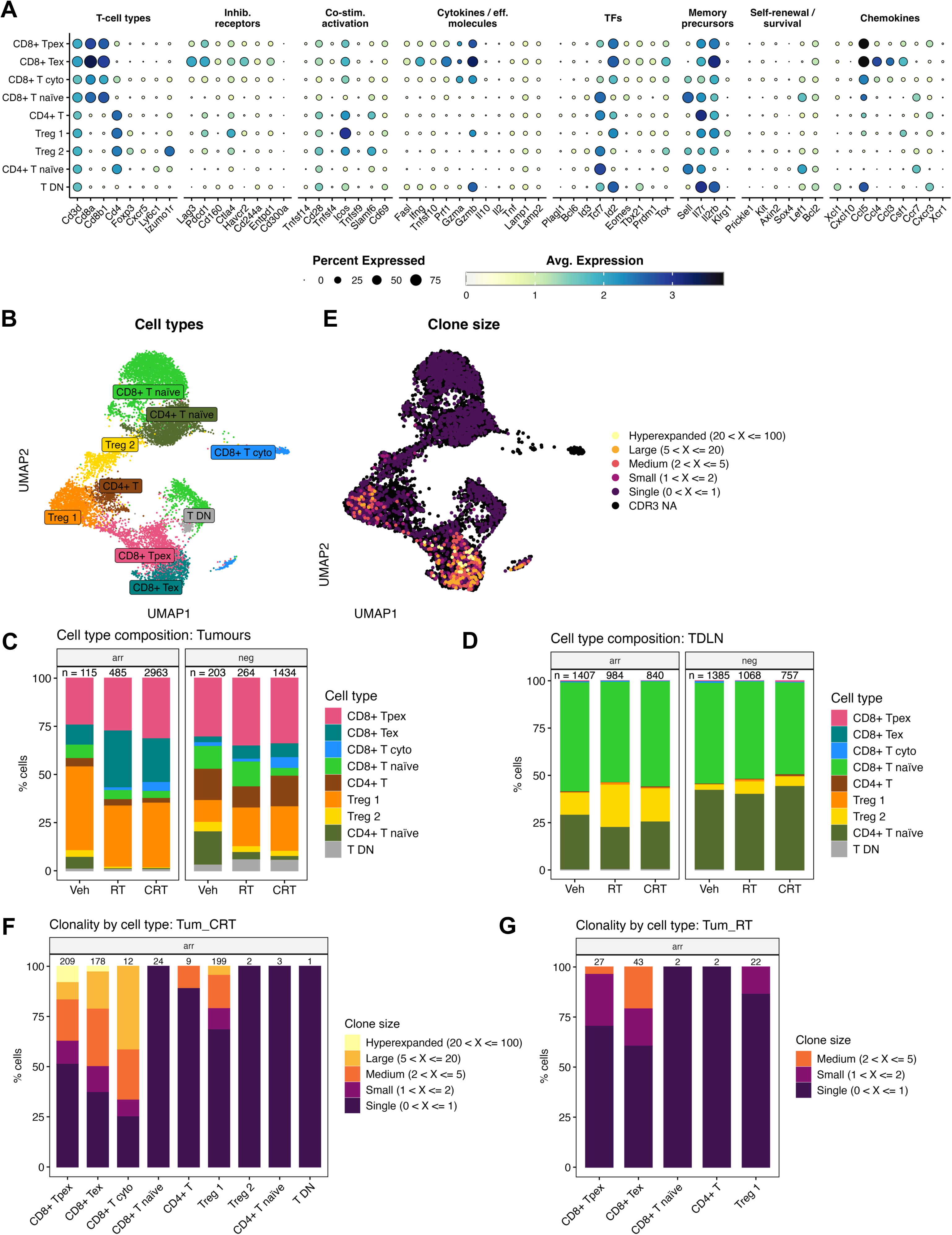
CRT increases the proportion and clonal expansion of precursor exhausted (T_PEX_) to terminally exhausted (T_EX_) CD8+ TILs in the arrested phase, compared to RT alone. Single-cell analysis whereby cells were subject to UMAP clustering based on transcriptomic profile (Vehicle n=12, RT n=12, CRT n=12, 1 experiment; only available for Tocky arrested [“arr”] and timer negative [“neg”] samples). **A.** Dot plot illustrating relative gene expression across different T-cell subgroups (each corresponding to a UMAP cluster) and gene categories. Dot colour intensity represents the average expression level, while dot size indicates the percentage of cells expressing the gene. **B.** Cell annotation of clusters by T-cell subgroups. **C-D.** Cell type composition by treatment group and Tocky stage “arr” or “neg” of tumour samples **(C)** and tumour-draining lymph nodes (TDLN) **(D)**. **E.** Clone size categories indicated by clone frequency within the total TCR repertoire: hyperexpanded (>20 and ≤100), large (>5 and ≤20), medium (>2 and ≤5), small (>1 and ≤2) and single (>0 and ≤1). Cells without CDR3 sequence information are denoted by “CDR3 NA”. **F-G:** Barplots illustrating percentage of cells within each clone size category by cell type cluster for “arr” Tum_CRT **(F)** and Tum_RT **(G)**. Abbreviations: Tpex; precursor exhausted T-cells, cyto, cytotoxic; Tex, terminally exhausted T-cells; Treg, regulatory T-cells; T DN, double negative T-cells; Inhib. Receptors, inhibitory receptors; Co-stim. activation; co-stimulatory/ activation; eff. molecules; effector molecules; TFs, transcription factors.

Consistent with AbSeq profiling, TILs were enriched for antigen-experienced populations, including CD8⁺ T_PEX_, CD8⁺ T_EX_, and Treg1 (Fig. 6C), whereas TDLNs were dominated by naïve CD8⁺ T-cells and Treg2 (Fig. 6D). Within the arrested TILs compartment, both RT and CRT increased the frequency of CD8⁺ T_EX_ compared to vehicle, with RT inducing a larger increase. Conversely, CRT preferentially expanded CD8⁺ T_PEX_ compared to RT (Fig. 6C). In the timer-negative compartment, both RT and CRT similarly enhanced the proportion of T_PEX and_ T_EX_ compared to vehicle. These findings suggest that, in cells with a history of antigen engagement, CRT skews differentiation toward progenitor-like states, while RT promotes more terminal T-cell exhaustion.

To explore transcriptional clonal dynamics, we integrated paired TCR VDJ and scRNA-seq profiles, mapping clonotypes onto UMAP-defined subsets (Fig. 6E). In total, 78.2% (1,410/1,803) of arrested and 81.8% (1,084/1,325) of timer-negative T-cells were clonotyped. Clonal expansion occurred mainly in “arrested” TILs rather than T-cells from TDLNs (Fig. S8). As shown in Fig. 6F, large and hyperexpanded clones were exclusively observed in CRT-treated TILs, predominantly within CD8⁺ T_PEX_ and, to a lesser extent, CD8⁺ T_EX_ and Treg1 subsets. Notably, the majority of hyperexpanded clones resided in the T_PEX_ population, whereas RT-induced expansion was restricted to smaller clones within T_EX_ clusters (Fig. 6G). Thus, CRT promotes robust clonal expansion of T_PEX_ cells, while RT favours expansion of terminally exhausted subsets. However, the presence of clonally expanded Treg1 population may contribute to an inhibitory TME, that limits the functional reinvigoration of CD8⁺ T-cells with anti-PD-1 (Fig. S5), despite CRT-induced expansion of T_PEX_ populations.

Overall, compared to vehicle, CRT reduced the proportion of T_EX_ CD8⁺ TILs and increased T_PEX_ populations, whereas RT drove the opposite trend. These results, supported by lineage and clonotype mapping, indicate that CRT preferentially maintains and/or expands progenitor-exhausted states, potentially preserving anti-tumour functionality compared to RT alone.

### Enhanced T-cell activation and altered exhaustion dynamics after CRT versus RT in human PBMCs

To provide proof-of-concept translational validation of CRT’s impact on the T-cell exhaustion trajectory versus RT, we analysed a small exploratory cohort of patients with LA-SCCHN receiving definitive RT (n=4) or CRT (n=8) as part of the INOVATE study (ISRCTN32335415). Peripheral blood mononuclear cells (PBMCs) were collected from patients receiving RT or CRT at baseline, week 3 of RT (RTW3), and 3 months post-treatment (RTFU), and analysed by flow cytometry. Over time, there was a trend toward a reduction in CD4⁺ Tconv and an increase in CD8⁺ T-cell frequencies related to CD45^+^ population, more pronounced in CRT than RT (Fig. 7A). At the 3-month follow-up, only CRT induced a significant fold change in CD8⁺ T-cells relative to baseline (Fig. 7B). CRT was also associated with increased Granzyme B⁺ CD8⁺ T-cells and CD56⁺ NK-like CD4⁺ Tconv (35–37) at follow-up, indicative of enhanced cytotoxic activity (Fig. 7C). In parallel, proliferating Ki67⁺ Treg were significantly reduced after CRT but not RT (Fig. 7D), while PD-1 expression on both CD4⁺ Tconv and CD8⁺ T-cells was increased after CRT (Fig. 7E). Examination of exhaustion marker trajectories revealed that both RT and CRT showed trends toward increasing putative T_EX_ CD8⁺ T-cells (TOX⁺ TCF-1⁻ PD-1^hi^) (38–40) during treatment (Fig. 7F). Notably, a significant increase in putative T_PEX_ cells (TCF-1⁺ TOX⁻ PD-1^int^) (5, 40, 41) emerged only after CRT at the 3-month follow-up (Fig. 7F). Hence, even with this limited number of patients, these data revealed longitudinal differences between RT and CRT in peripheral T-cell subset composition, cytotoxic activity, and exhaustion trajectories that mirror the data generated in the mouse model.

**Fig 7:**
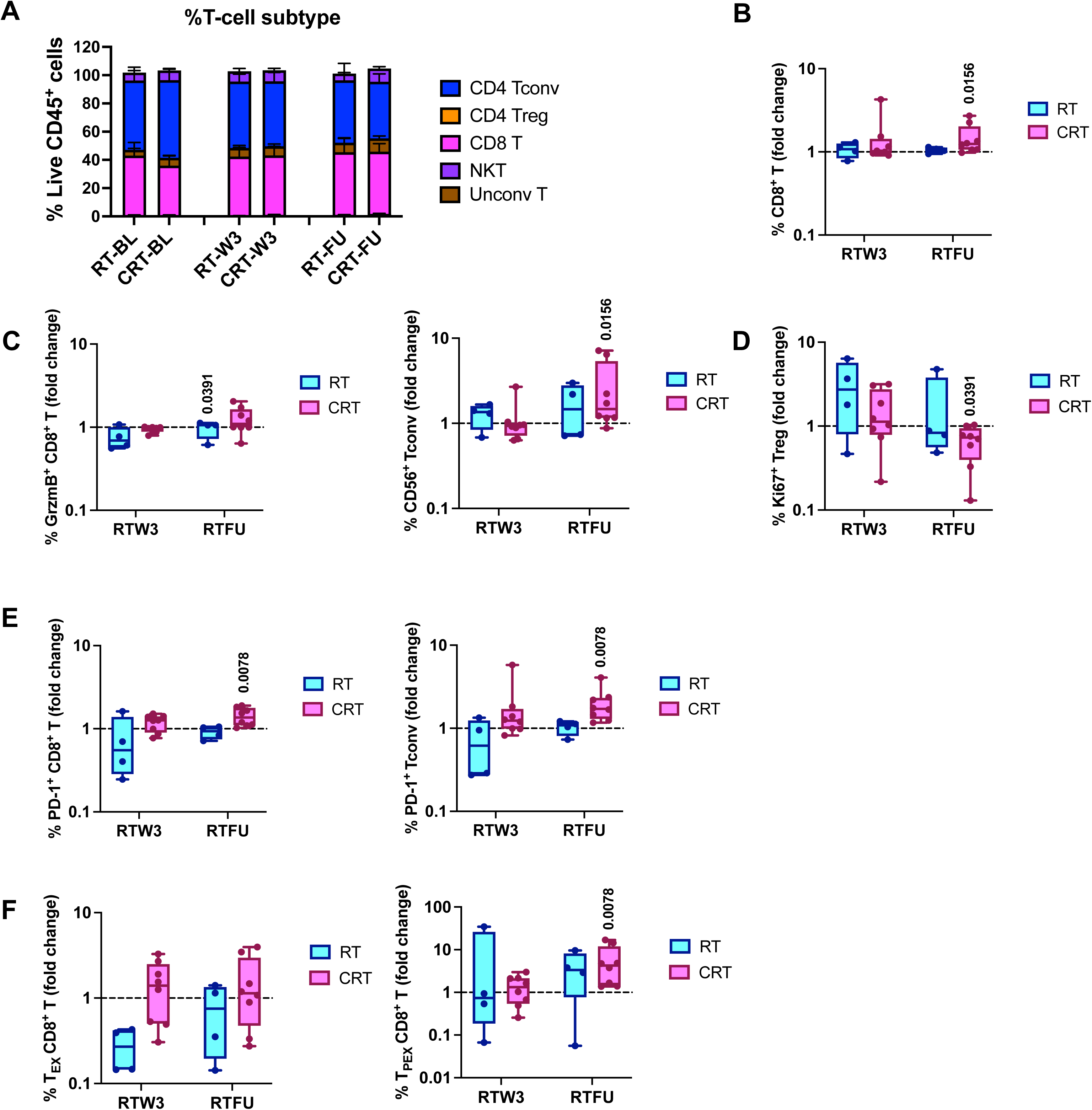
CRT versus RT differentially modulates T-cell subsets and exhaustion trajectories in human PBMCs. Peripheral blood mononuclear cells (PBMCs) were collected from patients with LA-SCCHN receiving definitive RT (n=4) or CRT (n=8) as part of the INOVATE study (ISRCTN32335415) at baseline (BL), week 3 of RT (RTW3), and 3 months post-treatment (RTFU; one RT patient at 6 weeks). Flow cytometry analysis of PBMCs was performed. **A.** Frequencies of T-cell subsets over time, comparing RT and CRT. **B-F.** Fold changes relative to baseline at RTW3 and RTFU shown as min-to-max box-and-whisker plots with all points displayed, for CD8⁺ T-cells **(B)**, Granzyme B⁺ (GrzmB^+^) CD8⁺ T-cells and CD56⁺ Tconv **(C)**, Ki67⁺ Tregs **(D)**, PD-1^+^ expression on CD8⁺ and Tconv **(E)**, terminally exhausted (T_EX_; TOX⁺ TCF-1⁻ PD-1^hi^) and progenitor exhausted (T_PEX_; TCF-1⁺ TOX⁻ PD-1^int^) CD8⁺ T-cell populations **(F)**. Statistical significance for fold change comparisons in panels B-F was assessed using the Wilcoxon signed-rank test; *p*<0.05.

## DISCUSSION

This study demonstrates that chemoradiotherapy (CRT) reshapes the intratumoural T-cell immunological landscape more favourably than radiotherapy (RT) alone in a relevant SCCHN tumour model (Fig S9). Functional CD8⁺ T-cell depletion experiments confirmed that tumour control under CRT, but not RT, depends on CD8⁺ T-cells, establishing their essential role in mediating CRT efficacy. Using the Nr4a3-Tocky reporter system combined with single-cell and clonotype analyses, we show that CRT enhances antigen engagement, promotes robust clonal expansion of CD8⁺ tumour-infiltrating lymphocytes (TILs), and biases differentiation toward a progenitor exhausted (T_PEX_), rather than terminally exhausted (T_EX_), fate. These immunological effects may be driven by the additive benefits of chemotherapy-induced immunogenic cell death and RT-mediated antigen release, which collectively broaden tumour antigen availability.

Both RT and CRT increased TIL activation and immune checkpoint expression; however, CRT induced a more clonally-focused CD8⁺ T-cell repertoire, including hyperexpanded clones, made up of only three paired TCR α/β CDR3 sequences (Fig. 4B), not observed with RT. These dominant clonotypes exhibited an effector-like phenotype with high CD44 and lower expression of inhibitory receptors (PD-1, LAG-3, TIGIT, CD38), suggesting functional persistence rather than terminal exhaustion. In contrast, RT-driven expansion was limited and enriched in T_EX_ subsets, a population associated with poor reinvigoration capacity. These observations are consistent with prior studies implicating T_PEX_ cells as key mediators of durable antitumour immunity and checkpoint blockade responsiveness (4, 6, 32).

Single-cell transcriptomics confirmed distinct T_EX_ and T_PEX_ states within TILs and revealed treatment-specific patterns. CRT enriched for T_PEX_ cells characterised by high *Tcf7*, intermediate *Pdcd1*, and low *Havcr2* expression, while RT promoted T_EX_ accumulation. Integration of clonotype and phenotype mapping revealed that large-to-hyperexpanded clones were restricted to CRT-treated tumours, primarily within T_PEX_ clusters, whereas RT-induced clonal expansion was sparse and largely confined to T_EX_ cells. These findings suggest that CRT not only broadens antigen engagement but also sustains clonal persistence in a less exhausted state, which may support combination immunotherapy. However, CRT clonally expanded Treg1, a Treg-like subset, with a suppressive transcriptional profile characterised by high PD-1, CTLA-4, ICOS, IL2RB, and ID2 expression which could counterbalance effector gains by limiting the functional reinvigoration of CD8⁺ T-cells. This may explain the limited clinical benefit of PD-1 blockade in this setting (consistent with our pre-clinical model data shown in Fig. S5), highlighting the need for additional combination strategies targeting regulatory populations such as anti-CD25 therapy. Interestingly, no shared clonotypes were detected between TDLNs and tumours, possibly due to limited sequencing depth, pooled sampling across animals, or the timing of tissue collection. However, shared clonotypes along the Tocky trajectory within TILs support antigen engagement as a driving force for clonal expansion within tumours.

From a translational perspective, these findings underscore the immunomodulatory potential of CRT beyond local tumour control. By preserving a pool of progenitor-like CD8⁺ T-cells and limiting terminal exhaustion, CRT may create a TME more able to contribute to systemic immune surveillance and amenable to reinvigoration by ICIs. Human PBMC analyses revealed that CRT induced persistently high putative T_EX_ CD8⁺ T-cells during treatment (week 3) and at 3 months post-treatment, while a significant increase in putative T_PEX_ cells emerged only at the later 3-month timepoint, and this was seen with CRT treatment only. This delayed T_PEX_ expansion may represent a recovery phase critical for effective antitumor immunity and responsiveness to ICIs. Concurrently, CRT was associated with increased cytotoxic potential, including higher Granzyme B⁺ CD8⁺ T-cells and CD56⁺ Tconv, as well as reduced proliferating Tregs, suggesting a shift toward a more active, systemic anti-tumour immune milieu. These data suggest that CRT modulates systemic lymphocyte immunity more favourably than RT during and after treatment. Importantly, these observations are supported by our group’s prior TCR sequencing work, which demonstrated that CRT promotes maintenance and expansion of pre-existing peripheral T-cell clonotypes associated with treatment response (9). However, as our analyses were performed in peripheral blood rather than tumour tissue, they fail to capture fully the intratumoral immune dynamics observed in our preclinical mouse models. The early and persistent dominance of systemic T_EX_ cells may also limit effective reinvigoration of T_PEX_ cells by PD-1-targeted therapies. These findings align with clinical trial data showing disappointing outcomes when ICIs are given concurrently with CRT in unresected LA-SCCHN (18–20). Our results highlight the importance of carefully timing ICI relative to CRT to capitalise on the potential window of progenitor T-cell expansion and improve therapeutic synergy.

Limitations of this study include modest single-cell RNA sequencing depth, which restricted the resolution of low-abundance transcripts and trajectory analysis. The exclusive analysis of sorted T-cells prevented broader immune cell interactions within the TME from being captured. Additionally, pooled tumour and TDLN samples from multiple mice may have masked individual-level clonal overlaps. The small number of patients in the translational cohort is also a limitation of this work. Future studies with longitudinal tumour sampling – both in preclinical models and from patients – together with deeper sequencing and paired tumour-TDLN analyses, could better clarify the dynamics of priming, trafficking, and differentiation in this setting.

Future clinical strategies should consider sequencing and biomarker-driven timing of immunotherapy. Rational combinatorial treatment strategies, that move beyond immune checkpoint inhibition (ICI) alone with CRT, could include Treg depletion (e.g. anti-CD25 antibodies) (42), targeting immunosuppressive myeloid populations (43), or radio-sensitising agents that enhance immunogenic cell death to further amplify antigen release and CD8⁺ T-cell priming (44). Such approaches may provide the additional immune support required to translate CRT-ICI synergy into meaningful clinical benefit.

In conclusion, our data demonstrate that the benefit of adding chemotherapy to radiotherapy extends beyond direct radiosensitisation of tumour cells. CRT remodels the TIL compartment by expanding antigen-engaged CD8⁺ T-cells, promoting clonal dominance, and sustaining progenitor-like states linked to durable antitumour immunity. These mechanistic insights into the T_PEX_/T_EX_ exhaustion trajectory provide a rationale for optimising the timing of CRT-based immunotherapy regimens in SCCHN, with future work focused on refining combination strategies that overcome residual immunosuppressive barriers.

## METHODS

### Mouse models of head and neck cancer

All procedures involving animals were approved by the Animal Ethics Committee at The Institute of Cancer Research in accordance with National Home Office Regulations under the Animals (Scientific Procedures) Act 1986. Wild-type or Nr4a3-Tocky C57BL/6 female mice (8-14 weeks) were housed under specific pathogen-free conditions with ad libitum food and water. The HPV-positive syngeneic mouse SCCHN mEER cell line was a gift from Dr Paola Vermeer (Sanford Research, USA) and HPV-negative C57BL/6 carcinogen-induced mouse oral cancer cell line MOC1 (45) were kindly provided by Ravindra Uppaluri (Dana-Farber Cancer Institute). Cells were cultured in complete DMEM (Gibco) supplemented with heat-inactivated 5/10% FBS, 1% L-glutamine and 0.5% penicillin-streptomycin. 1 × 10^6^ mEER cells or 4 × 10^6^ MOC1 cells were injected subcutaneously into the right flank under isoflurane anaesthesia. Tumour and weight measurements were taken twice weekly using calipers by an independent animal technician. Tumour volume (mm^3^) was calculated as length ξ width ξ height ξ 0.5236. For all studies, mice were randomised prior to treatment initiation for comparable tumour volumes between cohorts. Euthanasia was performed via cervical neck dislocation upon reaching humane endpoints, which included a maximum tumour diameter of 15 mm, ≥18% loss of body weight from peak weight, or clinical signs of distress.

### *In vivo* treatments

*In vivo* treatments were commenced at 14-18 days post-implantation for mEER or 28-35 days for MOC1, when tumour volumes reached approximately 30-60 mm^3^. For radiotherapy (RT), mice were anaesthetized with ketamine/xylazine and positioned prone under lead shielding with a 16 mm aperture over the tumour. mEER tumours received 8 Gy × 3 fractions and MOC1 tumours 6 Gy × 3 fractions on alternate days using an AGO 250 kV X-ray unit (dose rate: 1.62 Gy/min). Dosimetry was verified using a calibrated Farmer Chamber and Unidos-E Dosimeter (both PTW).

For chemoradiotherapy (CRT), cisplatin (7 mg/kg) was administered intraperitoneally 1 h before each RT fraction. Where indicated, immune checkpoint blockade was delivered via intraperitoneal anti–PD-1 (200 µg, clone RMP1-14) or isotype control twice weekly for up to six doses. CD8⁺ T-cell depletion was performed using anti-CD8α antibody (loading dose 400 µg, maintenance 200 µg twice weekly) or isotype control. Cisplatin (7 mg/kg) was administered by i.p. injection one hour prior to RT (if delivered). Where indicated, anti-PD-1 (200 µg, clone: RMP1-14, BioXCell) or rat IgG2a isotype control (clone: 2A3, BioXCell) were administered by i.p. injection twice-weekly for up to six doses. CD8^+^ T-cell depletion was performed using an anti-CD8α antibody (loading dose 400 µg, maintenance 200 µg twice weekly; clone: 2.43, BioXCell) or rat IgG2b isotype control (clone: 1-2, BioXCell) for up to 10 doses. Confirmation of *in vivo* CD8^+^ T-cell depletion using this regimen is outlined in Fig. S3.

### Histopathology

Mouse specimens were fixed in 10% neutral buffered formalin (Sigma-Aldrich, HT501128) for 24 hours at room temperature, after which they were transferred to PBS at 4°C. Paraffin embedding of fixed material and immunohistochemistry (IHC) staining was performed by the ICR Histopathology Core Facility. Anti-CD8⍺ (Abcam, clone: EPR21769) was used with heat induced epitope retrieval using Agilent Target Retrieval solution pH9 and Agilent rabbit EnVision reagent as the detection system. Slides were scanned and imaged using Hamamatsu Nanozoomer (Hamamatsu Photonics). Positive cell detection analysis on IHC slides was performed using QuPath v0.5.1.

### Flow cytometry analysis

Harvested mouse tumours were dissociated mechanically using scissors and enzymatically digested in PBS containing 0.5 mg/mL collagenase type 76 I-S (Sigma-Aldrich), 0.4 mg/mL Dispase II protease (Sigma-Aldrich), 0.2 mg/mL DNase I (Roche) and 4% Trypsin. Following digestion, samples were passed through a 70 μm cell strainer and washed with FACS buffer (PBS/2% FBS/5 mM EDTA). Lymph nodes were made into a single-cell suspension by passage through a 70 μm cell strainer. For some experiments, mice were bled 10-20 μL from the tail vein, and red blood cell lysis was performed by incubation in ACK lysis buffer (Thermo Fisher Scientific) for 2 min. Human cryopreserved PBMCs were thawed using CTL Anti-Aggregate Wash Supplement (Cellular Technology Limited) according to the manufacturer’s protocol. Samples were centrifuged at 300 g for 5 min at 4°C, washed and resuspended in FACS buffer for Fc blocking and staining. Cells were incubated 4 °C for 10 min with anti-mouse (BD Pharmingen) or anti-human CD16/CD32 (BD Pharmingen) prior to surface staining, 4 °C for 30 min. Dead cells were excluded by using the Fixable Viability Dye eFluor 780 (Thermo Fisher Scientific) and the complete list of antibodies used for staining in this study is provided in Table S1-S2. Intracellular staining was performed using the Foxp3/Transcription Factor Staining Buffer Set (Thermo Fisher Scientific) as per manufacturer’s protocol. All samples were acquired on a BD FACSymphony A5 Cell Analyser. Flow cytometry data was analysed using the FlowJo software (v10) and Tocky timer data was analysed using the “TockyAnalysis” package in R (v3.6.3). Gating strategies for different immune cell populations are detailed in Supplementary information (Fig. S10-S13).

### Single-cell RNA/TCR/Abseq profiling

Tumours and TDLNs from mEER tumour-bearing Nr4a3-Tocky mice were collected 10 days following the final dose of RT. Mice (n=12 per treatment group) received vehicle, RT, or CRT on day 14 post-tumour inoculation; one wild-type mouse was included for gating control. Tumours and TDLN were processed to form single-cell suspensions and pooled by treatment group and tissue type, generating six pooled samples. For tumour samples, samples were filtered and enriched for CD45^+^ cells with mouse CD45 (TIL) microbeads (Milentyi Biotec) according to manufacturer’s protocol with LS columns on the QuadroMACS magnet (Milentyi Biotec). Following this, cells were stained with AbSeq oligonucleotide-conjugated antibodies (Table S3) and sample tags for cellular indexing of transcriptomes and epitopes by sequencing (CITE-seq). Following flow cytometry staining, live TCRβ⁺ cells were sorted into four Tocky-defined populations (“new,” “persistent,” “arrested,” and “timer-negative”) using a BD FACSymphony S6 cell sorter, sorting strategies as detailed in Fig. S14. Sorted populations were processed on the BD Rhapsody system for single-cell capture, cDNA synthesis, and library preparation following the manufacturer’s protocol. Libraries for whole-transcriptome analysis (WTA), AbSeq, Sample Tags, and TCR sequencing were prepared and sequenced on Illumina NovaSeq platforms. Additive results from three sequencing runs (repeated due to technical issues), resulted in RNA libraries sequenced to a mean depth of 8,016 reads and 2,644 molecules per cell; AbSeq libraries reached a mean of 8,552 reads and 3,098 molecules per cell; and TCR libraries achieved a mean of 2,396 reads and 6.35 molecules per cell. Raw data was aligned and pre-processed using the BD Rhapsody pipeline and further filtered to retain cells with ≥10 detected proteins, ≥500 AbSeq unique molecular identifiers (UMIs), and mitochondrial content <20%. After quality control, at least 3,600 cells were retained in each of the “persistent”, “arrested”, and “timer-negative” populations. In contrast, relatively few “new” Tocky cells (∼200) passed QC, which limits the interpretability of analyses within this subset. Downstream analyses were performed in R (v4.3.3).

### Bulk RNA and TCR sequencing

Mouse tumours were harvested 14 days after the last radiation dose and snap frozen on dry ice. Tumour tissue was homogenised in 1 mL of Buffer RLT Plus (Qiagen) with 10 µL β-mercaptoethanol (Gibco) within CK28 hard tissue homogenising 2 mL tubes (Precellys) using the Precellys Evolution Homogeniser. RNA was extracted tumours using the RNeasy Plus Mini Kit with QIAshredder (Qiagen) according to manufacturer’s protocol. Library preparation and sequencing were performed by Genewiz, Germany. Library preparation was strand-specific with polyA selection, with 2×150bp sequencing at 20 million paired-end reads per sample performed on an Illumina NovaSeq. Salmon quant v1.10.1 (46) was used for transcript-level quantification against the GRCm39 mouse reference (Ensembl release 103). Transcript-level abundance estimates were aggregated to gene-level using tximport v1.34 R package (47). Differential gene expression analysis was performed using DESeq2 (48) (pseudogenes were excluded and only genes with more than 10 reads in at least 5 samples were kept in the analysis). Cell type signature genes were obtained from single-cell RNA sequencing data from oropharyngeal squamous cell carcinoma patients (49) and signature scores were calculated using the GSVA package (v2.0.4) (50) applied to counts normalised using variance stabilising transformation. Gene set enrichment analysis was performed using the clusterProfiler package (v4.14.4) (51).

### Human studies

Peripheral blood was collected from patients with LA-SCCHN treated with definitive RT (n=4) or CRT (n=8) as part of the INOVATE study (Investigation of Novel Plasma Human Papilloma Virus DNA Assay for Treatment Response Estimation in Head and Neck Cancer, ISRCTN32335415). All patients were aged ≥18 years with newly diagnosed HPV p16-positive T1-T2/N1-3 or T3-T4/N0-3 squamous cell carcinoma of the oropharynx undergoing radical treatment with RT or CRT. Samples were collected at baseline, week 3 of RT (RTW3), and at 3 months post-treatment (RTFU); one patient in the RT cohort had a follow-up collection at 6 weeks due to scheduling constraints. PBMCs were isolated using a density gradient medium (Lymphoprep, STEMCELL Technologies) and stored in liquid nitrogen until analysis.

### Statistical analysis

Statistical analyses were performed using GraphPad Prism version 10. Data are presented as means ± SEM and are derived from single or pooled results of two to three independent experiments. Area under the curve (AUC) was used to compare tumour growth between groups. Survival differences were assessed using the log-rank (Mantel-Cox) test. Parametric tests were applied only to data with normal distribution, confirmed via the Shapiro–Wilk test. For comparisons between two groups, either a two-tailed unpaired t-test or Mann–Whitney U test, or Wilcoxon signed-rank test (for fold change comparisons) was used, depending on data type and normality. For comparisons among multiple groups, either one-way ANOVA with Tukey’s multiple comparisons test or the Kruskal-Wallis test with Dunn’s correction was used, as appropriate. Significant outliers were excluded using the ROUT test (Q=1%). A *p*-value of <0.05 was considered statistically significant.

## DATA AVAILABILITY

The datasets generated will be deposited in a publicly available repository upon acceptance of the manuscript for publication.

## ACKNOWLEDGEMENTS

We are grateful to the team members of the Biological Service Unit, Flow Cytometry Facility and Breast Cancer Now Histopathology Core Facility at The Institute of Cancer Research for technical assistance. This work was supported by the Wellcome Trust, Oracle Cancer Trust, The Institute of Cancer Research/Royal Marsden Hospital (ICR/RM) Centre for Immunotherapy of Cancer, ICR/RM NIHR Biomedical Research Centre (IS-BRC-1215-20021), Cancer Research UK (CRUK) Head and Neck Programme Grant (DRCRPG-Nov22/100008) and ICR/RM CRUK RadNet Centre of Excellence (C7224/A28724).

## MATERIALS & CORRESPONDENCE

Correspondence and requests for materials should be addressed to Dr Charleen Chan Wah Hak, Division of Radiotherapy and Imaging, The Institute of Cancer Research, Chester Beatty Laboratories, 237 Fulham Road, London SW3 6JB.

## AUTHOR CONTRIBUTIONS

C.CWH., A.A.M., K.J.H., and A.R. contributed to the conception and design of the research as well as in writing the manuscript. C.CWH., A.R., E.C.P., V.R., L.C.H., M.G., E.S.A., S.F., I.D., A.B., J.N.K., H.B., C.M., M.P. performed the experiments and acquired data. A.P. provided bioinformatic analysis. C.CWH., A.R., E.C.P., A.P. and M.O. contributed to the analysis and interpretation of data. J.Y.L., P.N., H.N., M.C.U.C. and S.B. contributed to the collection and provision of clinical samples. M.O. provided the supply of Nr4a3-Tocky mice and Tocky data analysis bioinformatic pipelines. A.A.M. and K.J.H. provided funding for the project. A.A.M., K.J.H., and M.O. are the guarantors and joint senior authors of this work. All authors read and approved the final manuscript.

