## Supplementary information for "Addition of chemotherapy to radiotherapy promotes progenitor-exhausted CD8⁺ T-cell clonal dominance in head and neck cancer"

### Figure S1

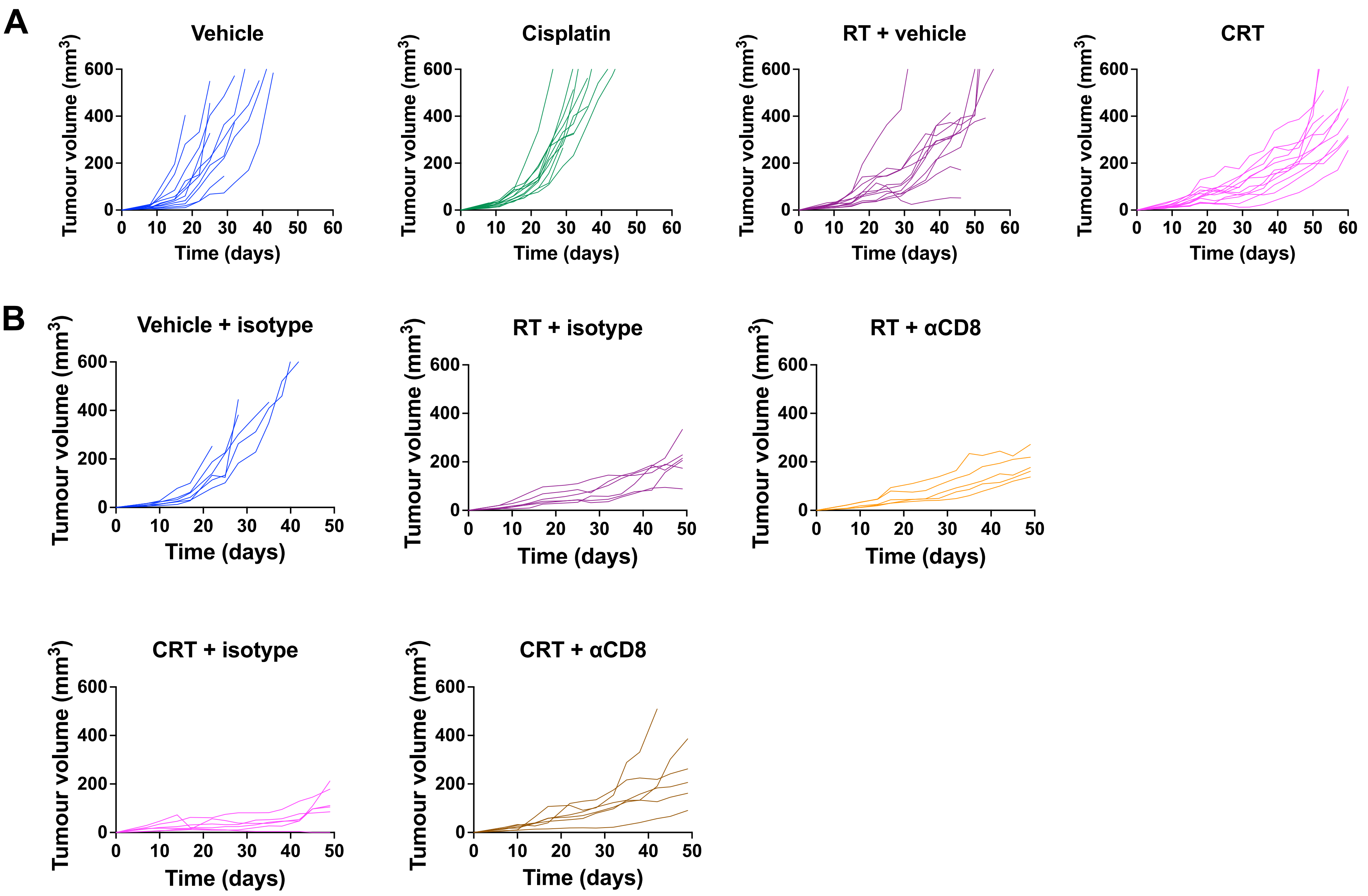
A
Vehicle
Cisplatin
RT + vehicle
CRT
Tumour volume (mm$^3$)
Time (days)
B
Vehicle + isotype
RT + isotype
RT + αCD8
CRT + isotype
CRT + αCD8

**Fig. S1: Individual tumour growth curves in mEER model receiving RT and CRT.**

**A.** CRT experiment (A-B: Vehicle n=11, Cisplatin n=11, RT + vehicle n=11, CRT n=12; 2 independent experiments). **B.** (C)RT plus $CD8^+$ T-cell depletion experiment (Vehicle + isotype n=6, RT + isotype n=6, RT + αCD8 n = 5, CRT + isotype n = 6, CRT + αCD8 n = 6; data from 1 experiment).

#### Figure S2

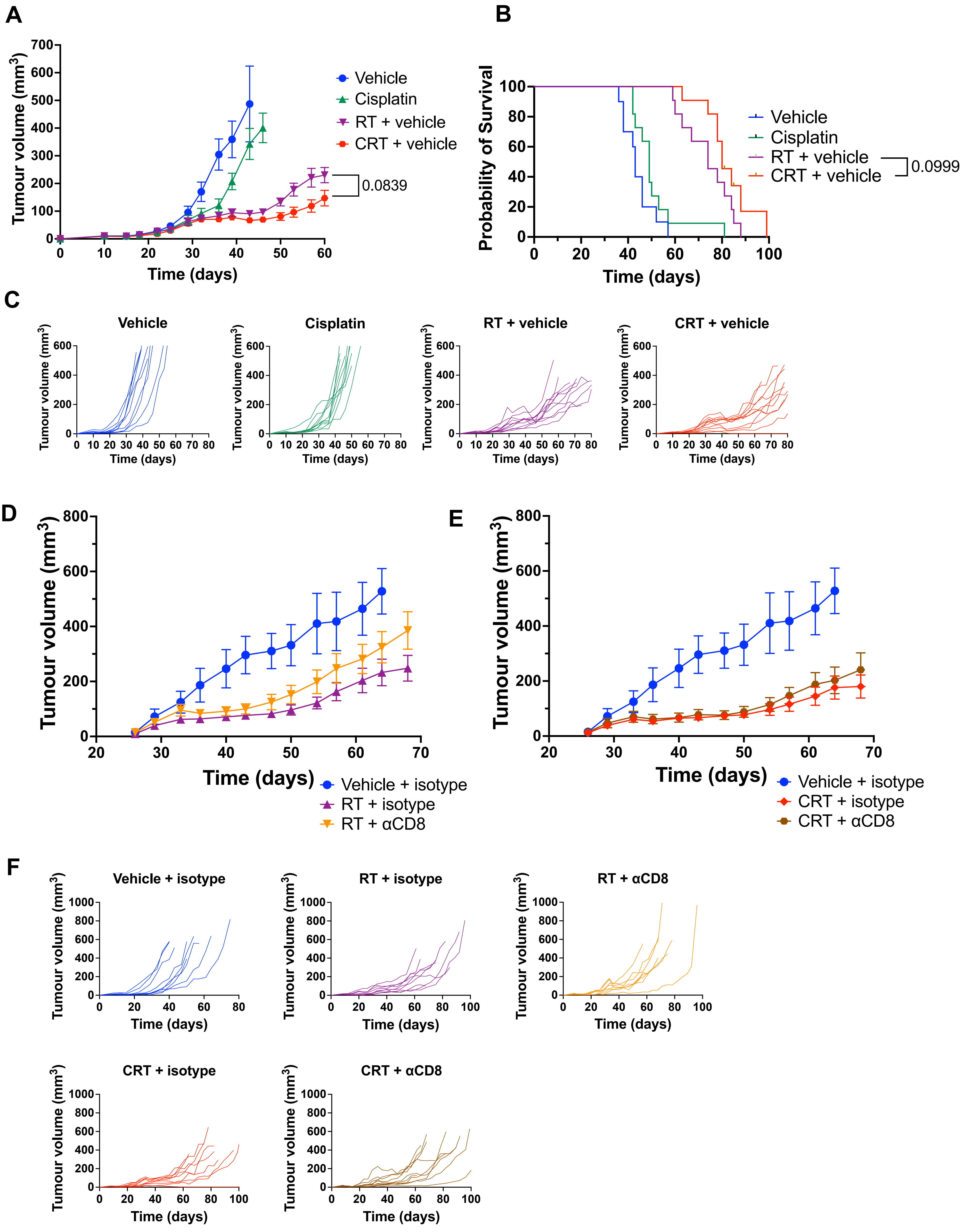
A
Tumour volume (mm3)
Time (days)
Vehicle
Cisplatin
RT + vehicle
CRT + vehicle
0.0839
B
Probability of Survival
Time (days)
Vehicle
Cisplatin
RT + vehicle
CRT + vehicle
0.0999
C
Vehicle
Cisplatin
RT + vehicle
CRT + vehicle
Tumour volume (mm3)
Time (days)
D
Tumour volume (mm3)
Time (days)
Vehicle + isotype
RT + isotype
RT + αCD8
E
Tumour volume (mm3)
Time (days)
Vehicle + isotype
CRT + isotype
CRT + αCD8
F
Vehicle + isotype
RT + isotype
RT + αCD8
CRT + isotype
CRT + αCD8
Tumour volume (mm3)
Time (days)

**Fig. S2: (C)RT treatment efficacy and CD8$^+$ T-cell dependence in the MOC1 model.**

**A-C:** (C)RT treatment regimen in the MOC1 model included RT 6 Gy × 3 alternate days and cisplatin/vehicle 5 mg/kg intraperitoneal (i.p.) injection once prior to the first fraction of RT (Vehicle n=10, Cisplatin n=10, RT + vehicle n=11, CRT + vehicle n= 11; 2 independent experiments). **(A)** Tumour growth curves across indicated treatment conditions. **(B)** Survival curves across indicated treatment conditions. **(C)** Individual tumour growth curves across indicated treatment conditions. **D-F**: *In vivo* CD8$^+$ T-cell depletion experiments (Vehicle + isotype n=9, RT + isotype n=8, RT + αCD8 n=8, CRT + isotype n=9, CRT + αCD8 n=10; 2 independent experiments). Average tumour growth curves in mice receiving αCD8 depleting antibody/ isotype with **(D)** RT or **(E)** CRT; and **(F)** individual growth curves. Results are means ± SEM and *n* denotes mice per group.

### Figure S3

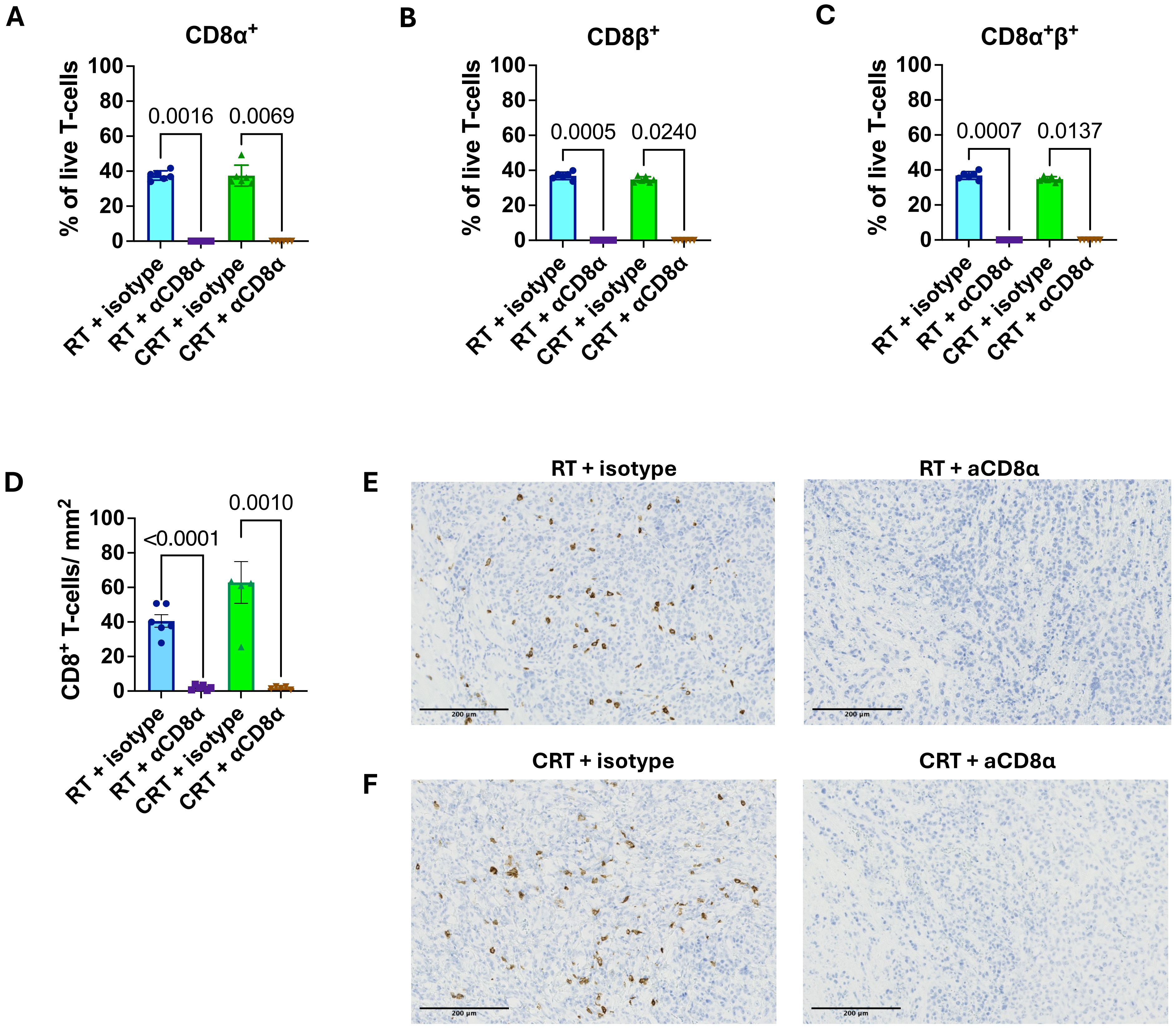

A
CD8α+
% of live T-cells
100
80
60
40
20
0
0.0016
0.0069
RT + isotype
RT + αCD8α
CRT + isotype
CRT + αCD8α
B
CD8β+
0.0005
0.0240
C
CD8α+β+
0.0007
0.0137
D
CD8+ T-cells/ mm2
<0.0001
0.0010
E
RT + isotype
RT + aCD8α
F
CRT + isotype
CRT + aCD8α
200 µm

**Fig. S3: Confirmation of peripheral blood and intratumoural CD8$^+$ T-depletion *in vivo* administration of a depleting anti-CD8α antibody.**

**A-C:** Peripheral blood from the tail vein of all mice from indicated treatment groups was collected two weeks following initiation of *in vivo* anti-CD8α (αCDα) depletion. Flow cytometry was performed to assess presence of **(A)** CD8α$^+$ **(B)** CD8β$^+$ and **(C)** CD8α$^+$β$^+$ T-cells. **D-F:** Tumours were collected at day 50 after tumour cell implantation and αCD8α immunohistochemistry (IHC) was performed. **(D)** Quantification of CD8$^+$ T-cells per mm$^2$ tumour area in αCD8α-treated versus isotype control-treated tumours. Representative IHC images were taken from **(E)** RT + isotype/αCD8α, **(F)** CRT + isotype/αCD8α. Sample sizes: RT + isotype n=6, RT + αCD8 n=6. CRT + isotype n=6, CRT + αCD8α n=6; panel D: RT + isotype n=6, RT + αCD8 n=6. CRT + isotype n=5, CRT + αCD8α n=5. Data are from one experiment.

### Figure S4

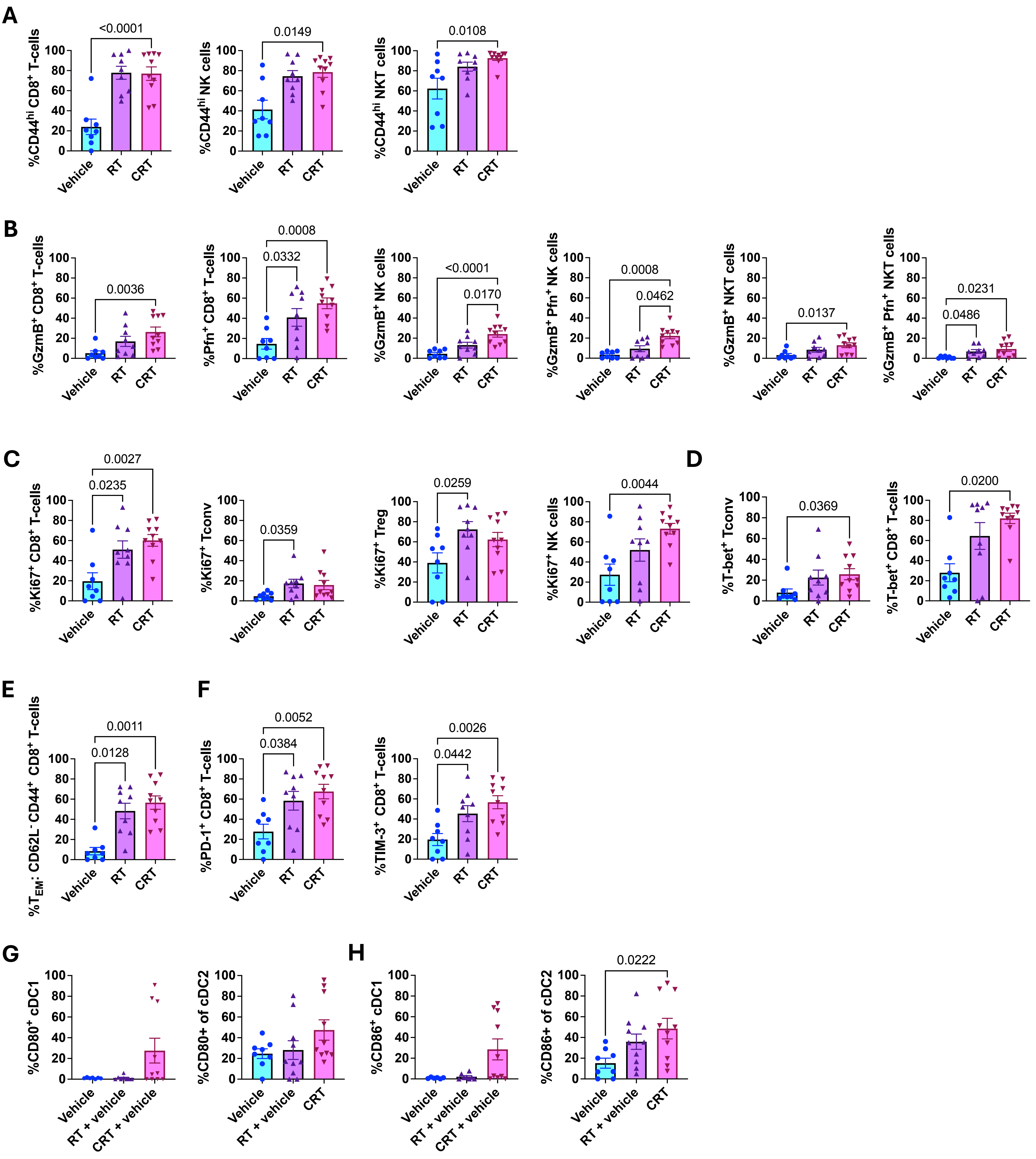
A
%CD44hi CD8+ T-cells
%CD44hi NK cells
%CD44hi NKT cells
<0.0001
0.0149
0.0108
B
%GzmB+ CD8+ T-cells
%Pfn+ CD8+ T-cells
%GzmB+ NK cells
%GzmB+ Pfn+ NK cells
%GzmB+ NKT cells
%GzmB+ Pfn+ NKT cells
0.0036
0.0008
0.0332
<0.0001
0.0170
0.0008
0.0462
0.0137
0.0231
0.0486
C
%Ki67+ CD8+ T-cells
%Ki67+ Tconv
%Ki67+ Treg
%Ki67+ NK cells
0.0027
0.0235
0.0359
0.0259
0.0044
D
%T-bet+ Tconv
%T-bet+ CD8+ T-cells
0.0369
0.0200
E
%TEM: CD62L- CD44+ CD8+ T-cells
0.0011
0.0128
F
%PD-1+ CD8+ T-cells
%TIM-3+ CD8+ T-cells
0.0052
0.0384
0.0026
0.0442
G
%CD80+ cDC1
%CD80+ of cDC2
H
%CD86+ cDC1
%CD86+ of cDC2
0.0222
0
20
40
60
80
100
Vehicle
RT
CRT
RT + vehicle
CRT + vehicle

**Fig. S4: Immunoprofiling via flow cytometry in mEER tumour-bearing mice.**

Mice received vehicle, RT or CRT as indicated and flow cytometric analysis was performed on tumours from each treatment group. **A.** Frequency (%) of $CD44^{hi}$ $CD8^+$ T-cells, NK cells, and NKT cells. **B.** Degranulation markers: frequency of Granzyme $B^+$ (GzmB) and $Perforin^+$ (Pfn) $CD8^+$ T-cells, $GzmB^+$ NK cells, $GzmB^+Pfn^+$ NK cells, $GzmB^+$ NKT cells, and $GzmB^+Pfn^+$ NKT cells. **C.** Frequency of $Ki67^+$ $CD8^+$ T-cells, Tconv, Treg and NK cells. **D.** Frequency of T-$bet^+$ Tconv and $CD8^+$ T-cells. **E.** Frequency of effector memory ($T_{em}$) $CD8^+$ T-cells. **F.** Frequency of PD-$1^+$ and TIM-$3^+$ $CD8^+$ T-cells. **G.** Frequency of $CD80^+$ conventional type 1 dendritic cells (cDC1) and conventional type 2 dendritic cells (cDC2). **H.** Frequency of CD86+ cDC1 and cDC2. Sample sizes: vehicle n=8, RT n=9, CRT n=10; for panels G-H: n=8, n=10, n=10. Data pooled from 2 independent experiments.

### Figure S5

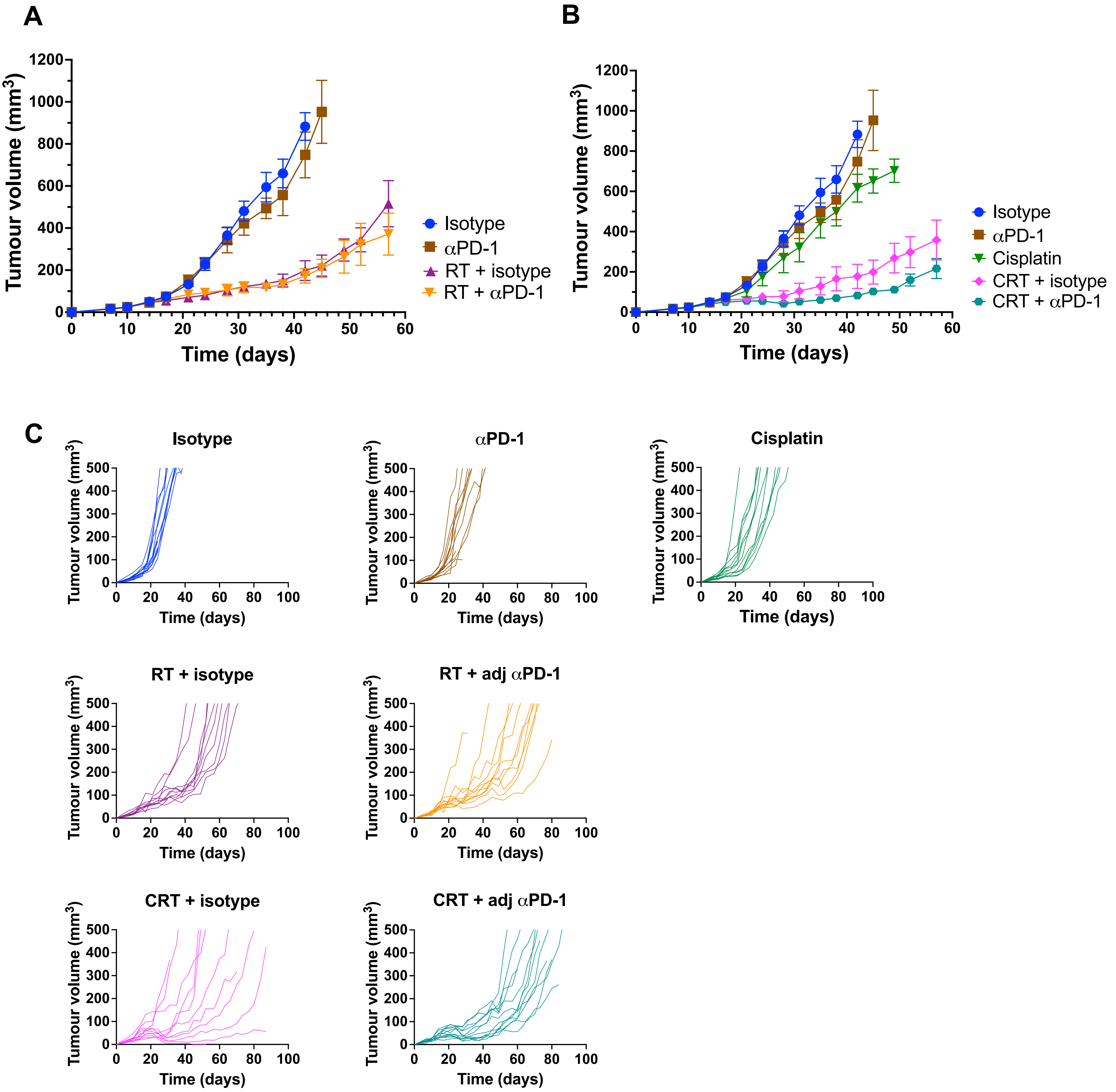
A
Tumour volume (mm³)
Time (days)
Isotype
αPD-1
RT + isotype
RT + αPD-1
B
Tumour volume (mm³)
Time (days)
Isotype
αPD-1
Cisplatin
CRT + isotype
CRT + αPD-1
C
Isotype
αPD-1
Cisplatin
RT + isotype
RT + adj αPD-1
CRT + isotype
CRT + adj αPD-1
Tumour volume (mm³)
Time (days)

**Fig. S5: *In vivo* effect of anti-PD-1 antibody on RT and CRT in the mEER model.**

**A-B:** Tumour growth curves across indicated treatment conditions: **(A)** RT plus anti-PD-1 experiment, (**B)** CRT plus anti-PD-1. **C.** Individual tumour growth curves (Isotype n=12, αPD-1 n=12, RT + isotype n=10, Cisplatin n=12, CRT + isotype n=11, RT + αPD-1 n=11, CRT + αPD-1 n=12; 2 independent experiments).

### Figure S6

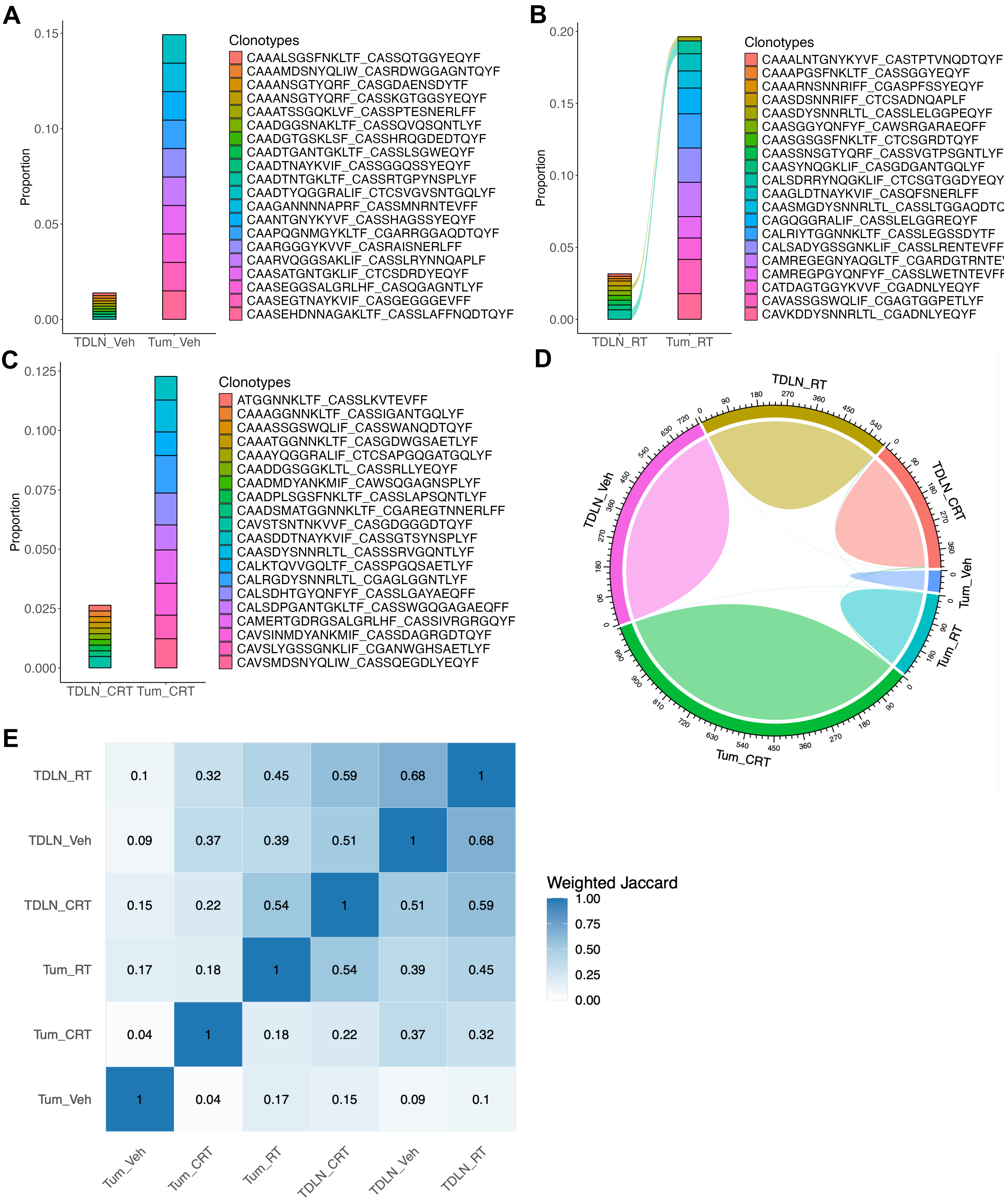
A
Proportion
0.15
0.10
0.05
0.00
TDLN_Veh
Tum_Veh
Clonotypes
CAAALSGSFNKLTF_CASSQTGGYEQYF
CAAAMDSNYQLIW_CASRDWGGAGNTQYF
CAAANSGTYQRF_CASGDAENSDYTF
CAAANSGTYQRF_CASSKGTGGSYEQYF
CAAATSSGQKLVF_CASSPTESNERLFF
CAADGGSNAKLTF_CASSQVQSQNTLYF
CAADGTGSKLSF_CASSHRQGDEDTQYF
CAADTGANTGKLTF_CASSLSGWEQYF
CAADTNAYKVIF_CASSGGQSSYEQYF
CAADTNTGKLTF_CASSRTGPYNSPLYF
CAADTYQGGRALIF_CTCSVGVSNTGQLYF
CAAGANNNNAPRF_CASSMNRNTEVFF
CAANTGNYKYVF_CASSHAGSSYEQYF
CAAPQGNMGYKLTF_CGARRGGAQDTQYF
CAARGGGYKVVF_CASRAISNERLFF
CAARVQGGSAKLIF_CASSLRYNNQAPLF
CAASATGNTGKLIF_CTCSDRDYEQYF
CAASEGGSALGRLHF_CASQGAGNTLYF
CAASEGTNAYKVIF_CASGEGGGEVFF
CAASEHDNNAGAKLTF_CASSLAFFNQDTQYF
B
Proportion
0.20
0.15
0.10
0.05
0.00
TDLN_RT
Tum_RT
Clonotypes
CAAALNTGNYKYVF_CASTPTVNQDTQYF
CAAAPGSFNKLTF_CASSGGYEQYF
CAAARNSNNRIFF_CGASPFSSYEQYF
CAASDSNNRIFF_CTCSADNQAPLF
CAASDYSNNRLTL_CASSLELGGPEQYF
CAASGGYQNFYF_CAWSRGARAEQFF
CAASGSGSFNKLTF_CTCSGRDTQYF
CAASSNSGTYQRF_CASSVGTPSGNTLYF
CAASYNQGKLIF_CASGDGANTGQLYF
CALSDRRYNQGKLIF_CTCSGTGGDYEQYF
CAAGLDTNAYKVIF_CASQFSNERLFF
CAASMGDYSNNRLTL_CASSLTGGAQDTQYF
CAGQGGRALIF_CASSLELGGREQYF
CALRIYTGGNNKLTF_CASSLEGSSDYTF
CALSADYGSSGNKLIF_CASSLRENTEVFF
CAMREGEGNYAQGLTF_CGARDGTRNTEVFF
CAMREGPGYQNFYF_CASSLWETNTEVFF
CATDAGTGGYKVVF_CGADNLYEQYF
CAVASSGSWQLIF_CGAGTGGPETLYF
CAVKDDYSNNRLTL_CGADNLYEQYF
C
Proportion
0.125
0.100
0.075
0.050
0.025
0.000
TDLN_CRT
Tum_CRT
Clonotypes
ATGGNNKLTF_CASSLKVTEVFF
CAAAGGNNKLTF_CASSIGANTGQLYF
CAAASSGSWQLIF_CASSWANQDTQYF
CAAATGGNNKLTF_CASGDWGSAETLYF
CAAAYQGGRALIF_CTCSAPGQGATGQLYF
CAADDGSGGKLTL_CASSRLLYEQYF
CAADMDYANKMIF_CAWSQGAGNSPLYF
CAADPLSGSFNKLTF_CASSLAPSQNTLYF
CAADSMATGGNNKLTF_CGAREGTNNERLFF
CAVSTSNTNKVVF_CASGDGGGDTQYF
CAASDDTNAYKVIF_CASSGTSYNSPLYF
CAASDYSNNRLTL_CASSSRVGQNTLYF
CALKTQVVGQLTF_CASSPGQSAETLYF
CALRGDYSNNRLTL_CGAGLGGNTLYF
CALSDHTGYQNFYF_CASSLGAYAEQFF
CALSDPGANTGKLTF_CASSWGQGAGAEQFF
CAMERTGDRGSALGRLHF_CASSIVRGRGQYF
CAVSINMDYANKMIF_CASSDAGRGDTQYF
CAVSLYGSSGNKLIF_CGANWGHSAETLYF
CAVSMDSNYQLIW_CASSQEGDLYEQYF
D
TDLN_RT
TDLN_CRT
Tum_Veh
Tum_RT
Tum_CRT
TDLN_Veh
E
TDLN_RT 0.1 0.32 0.45 0.59 0.68 1
TDLN_Veh 0.09 0.37 0.39 0.51 1 0.68
TDLN_CRT 0.15 0.22 0.54 1 0.51 0.59
Tum_RT 0.17 0.18 1 0.54 0.39 0.45
Tum_CRT 0.04 1 0.18 0.22 0.37 0.32
Tum_Veh 1 0.04 0.17 0.15 0.09 0.1
Tum_Veh
Tum_CRT
Tum_RT
TDLN_CRT
TDLN_Veh
TDLN_RT
Weighted Jaccard
1.00
0.75
0.50
0.25
0.00

**Fig. S6: Lack of shared clonotypes between tumour and TDLN sample types of the same treatment group.**

Single-cell TCR-seq analysis was performed on sorted tumour-infiltrating lymphocytes (TILs) from tumours (Tum) or TDLN harvested from mEER-bearing mice treated with vehicle (Veh), RT or CRT (Vehicle n=12, RT n=12, CRT n=12, 1 experiment). **A-C.** Alluvial plots tracking the top 10 clonotypes between sample types (Tumour [Tum] and TDLN) of the same treatment group: Vehicle **(A)**, RT **(B)**, CRT **(C)**. **D.** Chord diagram illustrating lack of shared clonotypes between all samples. **E.** Frequency-weighted 3-mer Jaccard similarity heatmap, visualising the degree of TCR repertoire similarity between samples.

### Figure S7

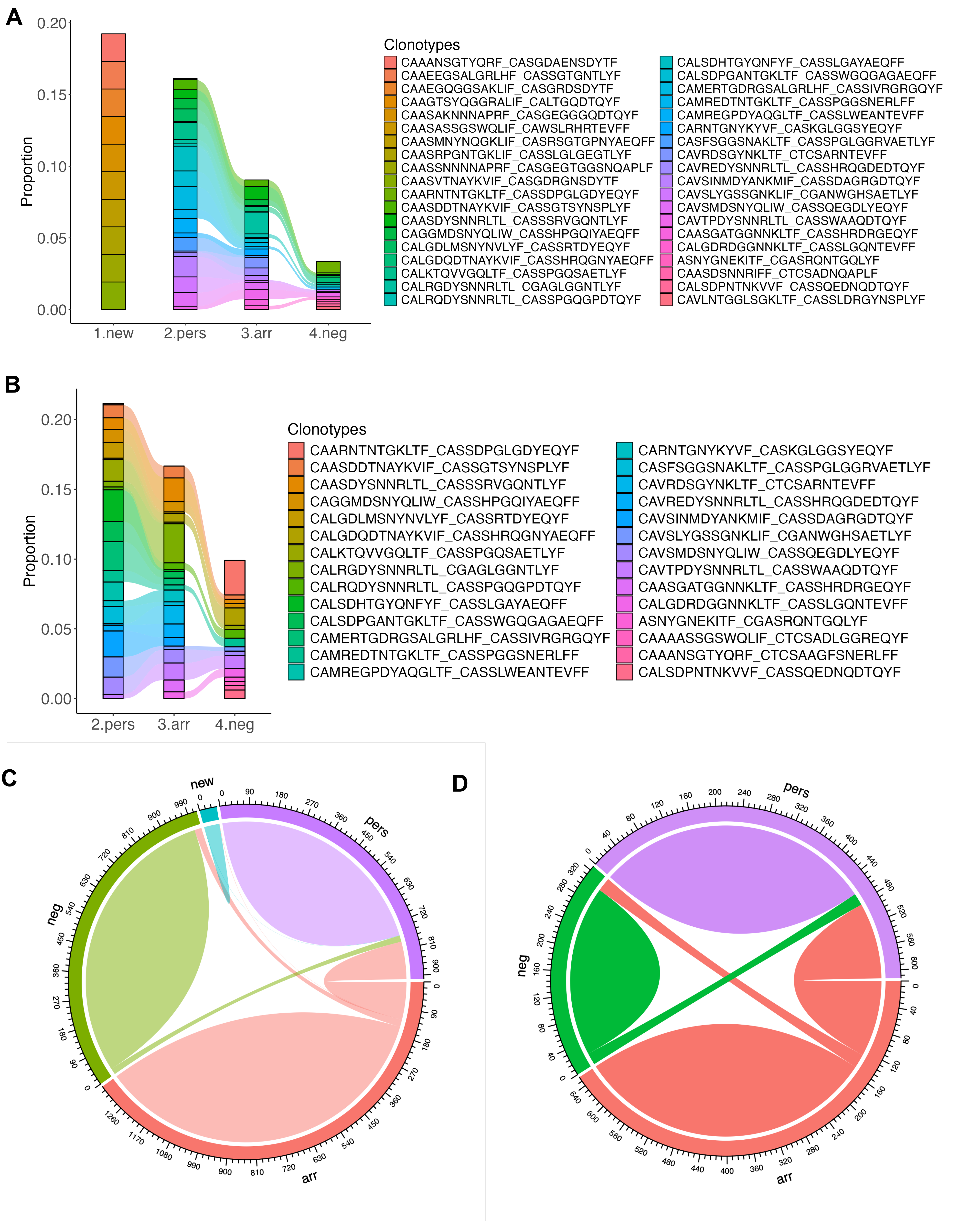
A
Proportion
0.20
0.15
0.10
0.05
0.00
1.new
2.pers
3.arr
4.neg
Clonotypes
CAAANSGTYQRF_CASGDAENSDYTF
CAAEEGSALGRLHF_CASSGTGNTLYF
CAAEGQGGSAKLIF_CASGRDSDYTF
CAAGTSYQGGRALIF_CALTGQDTQYF
CAASAKNNNAPRF_CASGEGGGQDTQYF
CAASASSGSWQLIF_CAWSLRHRTEVFF
CAASMNYNQGKLIF_CASRSGTGPNYAEQFF
CAASRPGNTGKLIF_CASSLGLGEGTLYF
CAASSNNNNAPRF_CASGEGTGGSNQAPLF
CAASVTNAYKVIF_CASGDRGNSDYTF
CAARNTNTGKLTF_CASSDPGLGDYEQYF
CAASDDTNAYKVIF_CASSGTSYNSPLYF
CAASDYSNNRLTL_CASSSRVGQNTLYF
CAGGMDSNYQLIW_CASSHPGQIYAEQFF
CALGDLMSNYNVLYF_CASSRTDYEQYF
CALGDQDTNAYKVIF_CASSHRQGNYAEQFF
CALKTQVVGQLTF_CASSPGQSAETLYF
CALRGDYSNNRLTL_CGAGLGGNTLYF
CALRQDYSNNRLTL_CASSPGQGPDTQYF
CALSDHTGYQNFYF_CASSLGAYAEQFF
CALSDPGANTGKLTF_CASSWGQGAGAEQFF
CAMERTGDRGSALGRLHF_CASSIVRGRGQYF
CAMREDTNTGKLTF_CASSPGGSNERLFF
CAMREGPDYAQGLTF_CASSLWEANTEVFF
CARNTGNYKYVF_CASKGLGGSYEQYF
CASFSGGSNAKLTF_CASSPGLGGRVAETLYF
CAVRDSGYNKLTF_CTCSARNTEVFF
CAVREDYSNNRLTL_CASSHRQGDEDTQYF
CAVSINMDYANKMIF_CASSDAGRGDTQYF
CAVSLYGSSGNKLIF_CGANWGHSAETLYF
CAVSMDSNYQLIW_CASSQEGDLYEQYF
CAVTPDYSNNRLTL_CASSWAAQDTQYF
CAASGATGGNNKLTF_CASSHRDRGEQYF
CALGDRDGGNNKLTF_CASSLGQNTEVFF
ASNYGNEKITF_CGASRQNTGQLYF
CAASDSNNRIFF_CTCSADNQAPLF
CALSDPNTNKVVF_CASSQEDNQDTQYF
CAVLNTGGLSGKLTF_CASSLDRGYNSPLYF
B
Proportion
0.20
0.15
0.10
0.05
0.00
2.pers
3.arr
4.neg
Clonotypes
CAARNTNTGKLTF_CASSDPGLGDYEQYF
CAASDDTNAYKVIF_CASSGTSYNSPLYF
CAASDYSNNRLTL_CASSSRVGQNTLYF
CAGGMDSNYQLIW_CASSHPGQIYAEQFF
CALGDLMSNYNVLYF_CASSRTDYEQYF
CALGDQDTNAYKVIF_CASSHRQGNYAEQFF
CALKTQVVGQLTF_CASSPGQSAETLYF
CALRGDYSNNRLTL_CGAGLGGNTLYF
CALRQDYSNNRLTL_CASSPGQGPDTQYF
CALSDHTGYQNFYF_CASSLGAYAEQFF
CALSDPGANTGKLTF_CASSWGQGAGAEQFF
CAMERTGDRGSALGRLHF_CASSIVRGRGQYF
CAMREDTNTGKLTF_CASSPGGSNERLFF
CAMREGPDYAQGLTF_CASSLWEANTEVFF
CARNTGNYKYVF_CASKGLGGSYEQYF
CASFSGGSNAKLTF_CASSPGLGGRVAETLYF
CAVRDSGYNKLTF_CTCSARNTEVFF
CAVREDYSNNRLTL_CASSHRQGDEDTQYF
CAVSINMDYANKMIF_CASSDAGRGDTQYF
CAVSLYGSSGNKLIF_CGANWGHSAETLYF
CAVSMDSNYQLIW_CASSQEGDLYEQYF
CAVTPDYSNNRLTL_CASSWAAQDTQYF
CAASGATGGNNKLTF_CASSHRDRGEQYF
CALGDRDGGNNKLTF_CASSLGQNTEVFF
ASNYGNEKITF_CGASRQNTGQLYF
CAAAASSGSWQLIF_CTCSADLGGREQYF
CAAANSGTYQRF_CTCSAAGFSNERLFF
CALSDPNTNKVVF_CASSQEDNQDTQYF
C
new
pers
arr
neg
D
pers
arr
neg

**Fig. S7: Shared clonotypes are seen to persist according to the Tocky trajectory of antigen engagement.**

Single-cell TCR-seq analysis was performed on sorted tumour-infiltrating lymphocytes (TILs) from tumours (Tum) or TDLN harvested from mEER-bearing mice treated with vehicle (Veh), RT or CRT (Vehicle n=12, RT n=12, CRT n=12, 1 experiment). **A-B.** Alluvial plots tracking the top 10 clonotypes across the different Tocky populations: “new”, “persistent” (pers), “arrested” (arr) and “timer negative” (neg), for all samples (Tumour and TDLN) **(A)** and Tumour samples alone **(B). C-D.** Chord diagrams showing shared clonotypes across different Tocky populations, for all samples (Tumour and TDLN) **(C)** and Tumour samples alone **(D)**.

### Figure S8

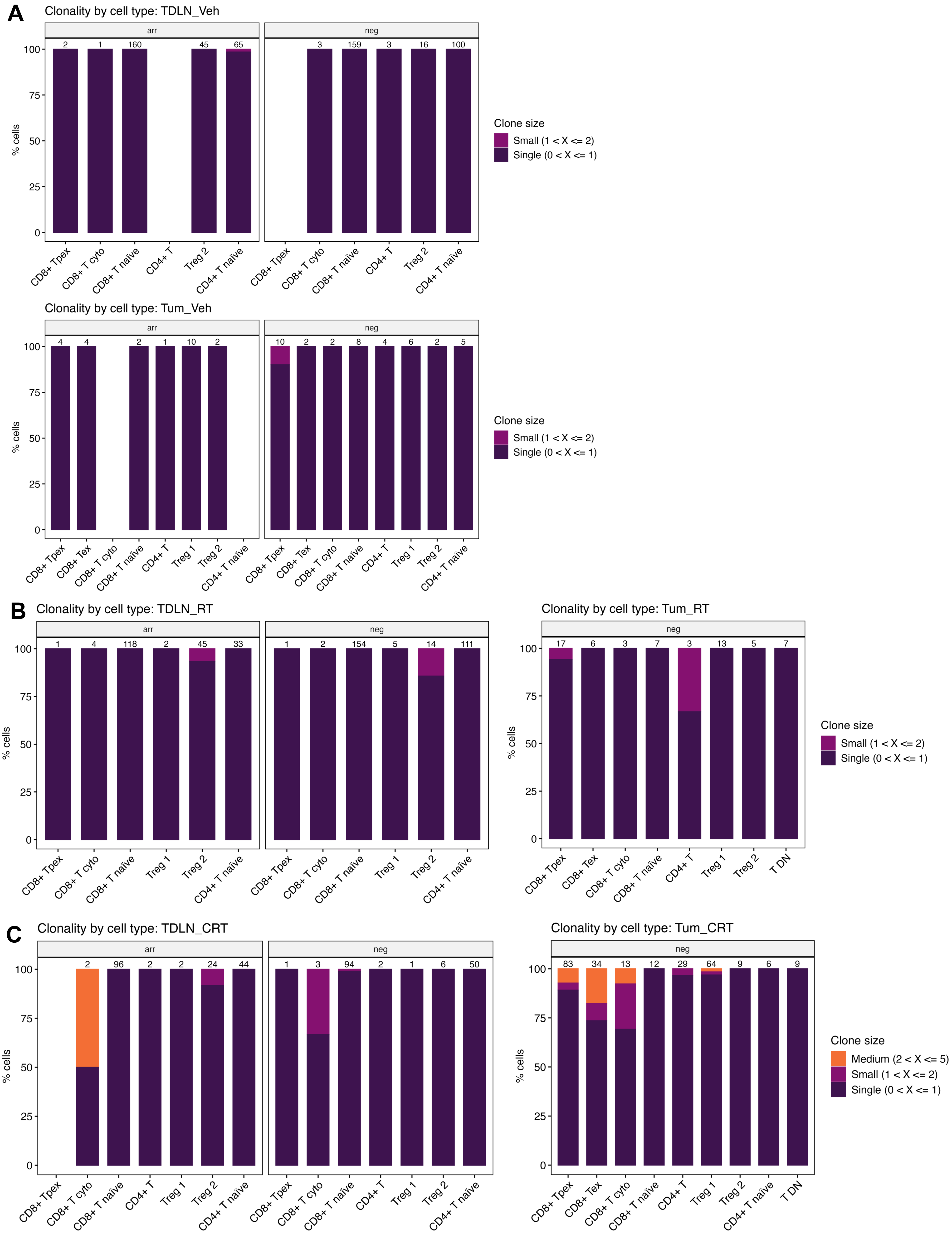

A
Clonality by cell type: TDLN_Veh
arr
neg
2
1
160
45
65
3
159
3
16
100
% cells
100
75
50
25
0
CD8+ Tpex
CD8+ T cyto
CD8+ T naïve
CD4+ T
Treg 2
CD4+ T naïve
Clone size
Small (1 < X <= 2)
Single (0 < X <= 1)
Clonality by cell type: Tum_Veh
4
4
2
1
10
2
10
2
2
8
4
6
2
5
CD8+ Tex
Treg 1
B
Clonality by cell type: TDLN_RT
1
4
118
2
45
33
1
2
154
5
14
111
Clonality by cell type: Tum_RT
17
6
3
7
3
13
5
7
T DN
C
Clonality by cell type: TDLN_CRT
2
96
2
2
24
44
1
3
94
2
1
6
50
Clonality by cell type: Tum_CRT
83
34
13
12
29
64
9
6
9
Medium (2 < X <= 5)

**Fig. S8: Clonality group by sample type, cell type and Tocky loci.**

Single-cell analysis whereby cells were subject to UMAP clustering based on transcriptomic profile (Vehicle n=12, RT n=12, CRT n=12, 1 experiment; only available for Tocky arrested ["arr"] and timer negative ["neg"] samples). **A-C:** Barplots illustrating percentage of cells within each clone size category by cell type cluster for **(A)** Vehicle-treated TDLN (TDLN_Veh) and tumours (Tum_Veh), **(B)** RT-treated TDLN (TDLN_RT) and tumours (Tum_RT) [Tum_RT "arr" data displayed in Fig. 6G] and **(C)** CRT-treated TDLN (TDLN_CRT) and tumours (Tum_CRT) [Tum_CRT "arr" data displayed in Fig. 6F]. Abbreviations: Tpex; precursor exhausted T-cells, cyto, cytotoxic; Tex, terminally exhausted T-cells; Treg, regulatory T-cells; T DN, double negative T-cells; Inhib. Receptors, inhibitory receptors; Co-stim. activation; co-stimulatory/ activation; eff. molecules; effector molecules; TFs, transcription factors; cyto, cytotoxic; Treg, regulatory T-cells.

**Figure S9**

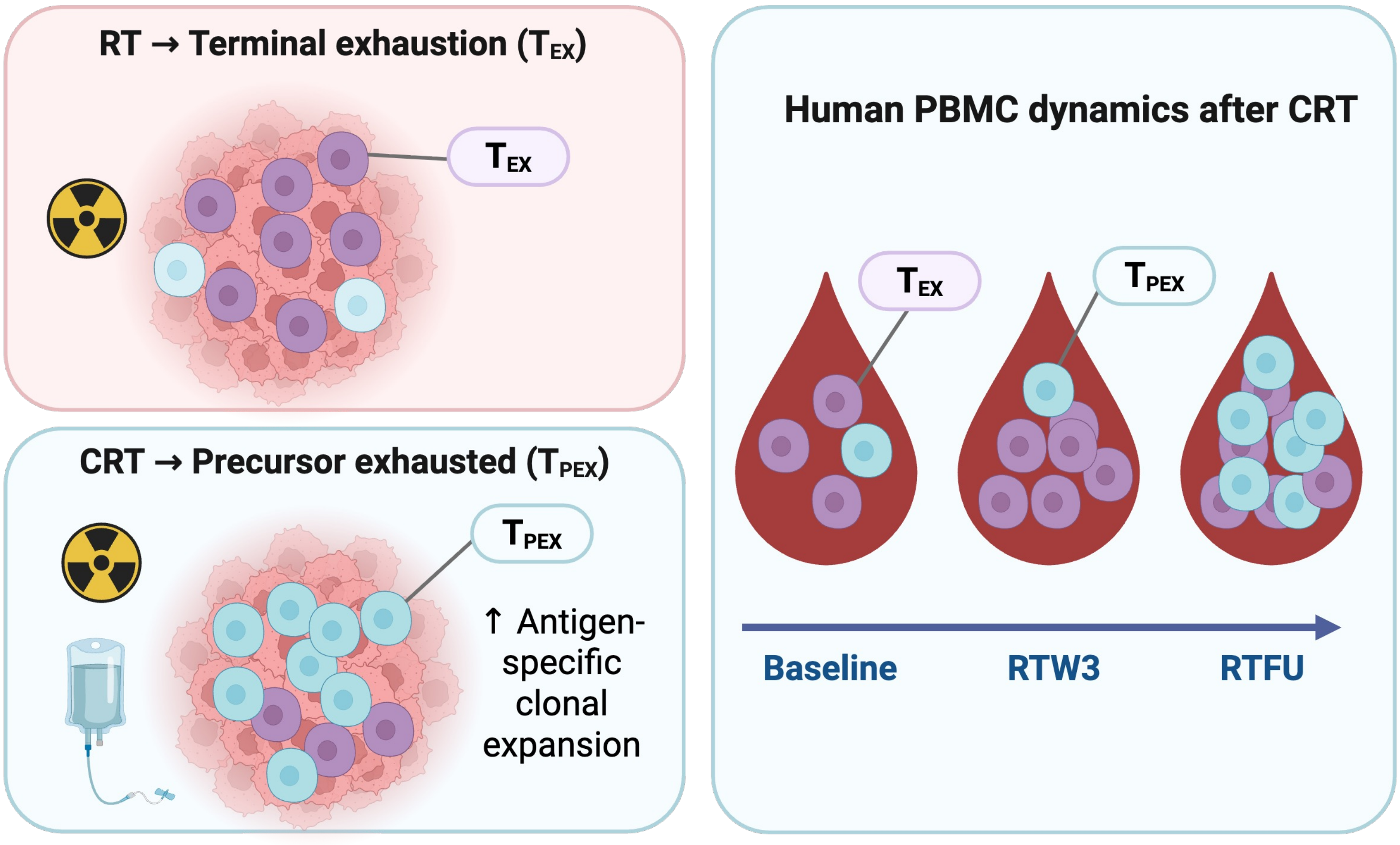

RT → Terminal exhaustion ($T_{EX}$)
$T_{EX}$
CRT → Precursor exhausted ($T_{PEX}$)
$T_{PEX}$
↑ Antigen-specific clonal expansion
Human PBMC dynamics after CRT
$T_{EX}$
$T_{PEX}$
Baseline
RTW3
RTFU

**Fig. S9: Distinct CD8$^+$ T-cell remodeling by chemoradiotherapy versus radiotherapy.** Graphical abstract illustrating how CRT, compared to RT alone, differentially shapes CD8$^+$ T-cell states in pre-clinical models of head and neck squamous cell carcinoma. CRT drives progenitor-exhausted ($T_{PEX}$) clonal expansion within tumours, while RT favours terminal exhaustion ($T_{EX}$). Human PBMC analyses following CRT show delayed recovery of $T_{PEX}$ cells 3 months post-RT (RTFU), highlighting a potential window for immunotherapy. RTW3: during week 3 of RT.

### Figure S10

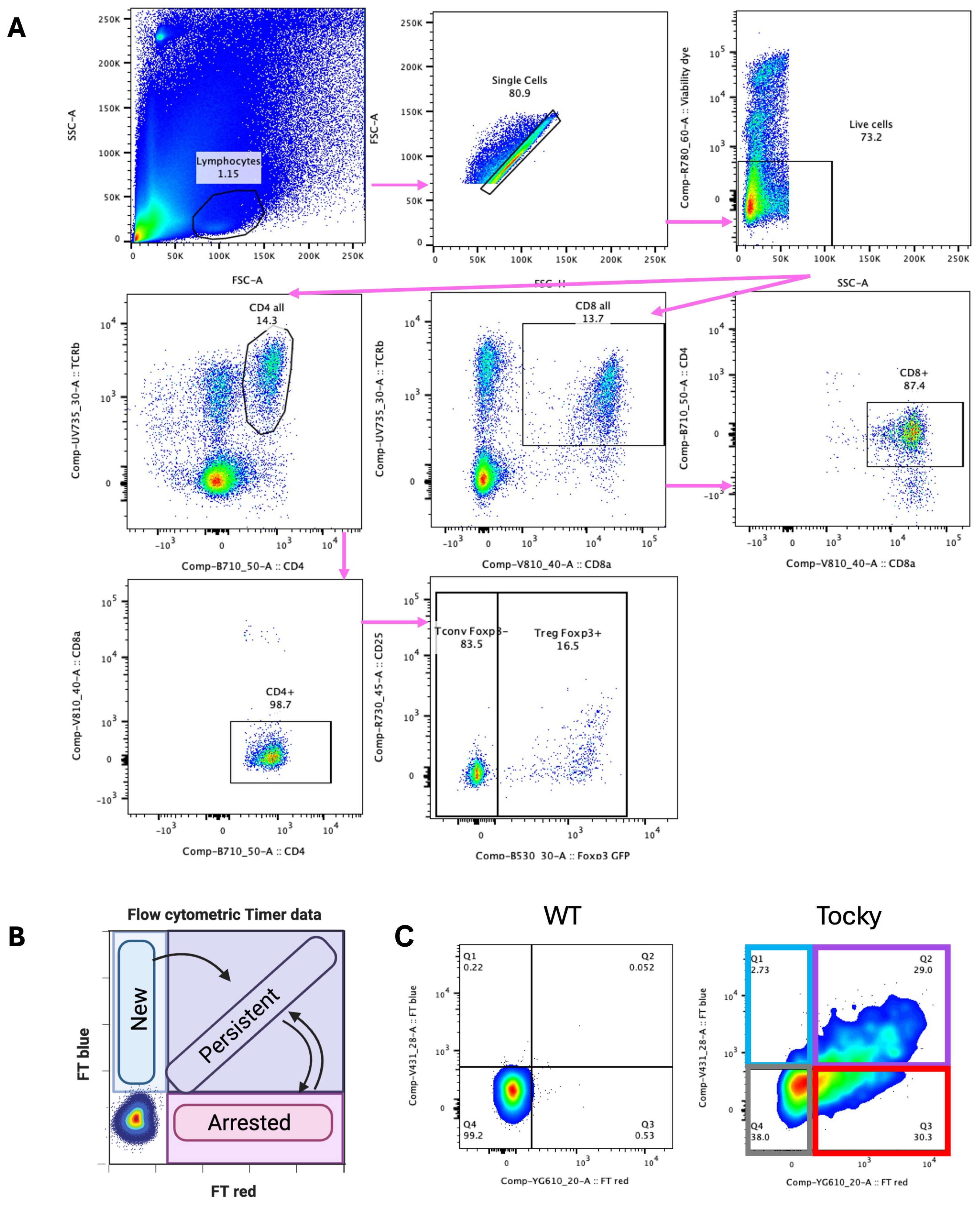
A
Lymphocytes
1.15
SSC-A
FSC-A
Single Cells
80.9
FSC-H
Live cells
73.2
Comp-R780_60-A :: Viability dye
CD4 all
14.3
Comp-UV735_30-A :: TCRb
Comp-B710_50-A :: CD4
CD8 all
13.7
Comp-V810_40-A :: CD8a
CD8+
87.4
CD4+
98.7
Tconv Foxp3-
83.5
Treg Foxp3+
16.5
Comp-R730_45-A :: CD25
Comp-B530_30-A :: Foxp3 GFP
B
Flow cytometric Timer data
New
Persistent
Arrested
FT blue
FT red
C
WT
Q1
0.22
Q2
0.052
Q4
99.2
Q3
0.53
Comp-V431_28-A :: FT blue
Comp-YG610_20-A :: FT red
Tocky
Q1
2.73
Q2
29.0
Q4
38.0
Q3
30.3

**Fig. S10: Nr4a3-Tocky lymphocyte panel gating strategy.**

The Nr4a3-Tocky reporter system enables *in vivo* analysis of T-cell activation dynamics via TCR engagement. Upon TCR stimulation and Nr4a3 transcription, an unstable fluorescent timer protein is initially expressed as a blue fluorophore, which matures into a red fluorophore over several hours. Newly engaged T-cells are blue-only, persistently engaged T-cells express both blue and red, arrested cells are red-only, and timer-negative cells lack both signals. **A.** All populations were gated from live single cells: conventional $CD4^+$ T-cells (Tconv), $TCR\beta^+CD4^+Foxp3^-$; regulatory T-cells (Treg), $TCR\beta^+CD4^+Foxp3^+$; $CD8^+$ T-cells, $TCR\beta^+CD8^+$. **B.** Schematic of flow cytometry timer data gating with fluorescent timer (FT) blue on the *y*-axis and FT red on the *x*-axis, classifying T-cells into "new" (FT $blue^+$ FT $red^-$), "persistent" (FT $blue^+$ FT $red^+$), "arrested" (FT $blue^-$ FT $red^+$), and "timer negative" (FT $blue^-$ FT $red^-$) populations. **C.** Example flow cytometry gating of Tocky FT populations set on the corresponding T-cell population in a wild-type (WT) sample, applied to a Tocky sample to define the "new" (blue), "persistent" (purple), "arrested" (red), and "timer negative" (grey) subsets.

### Figure S11

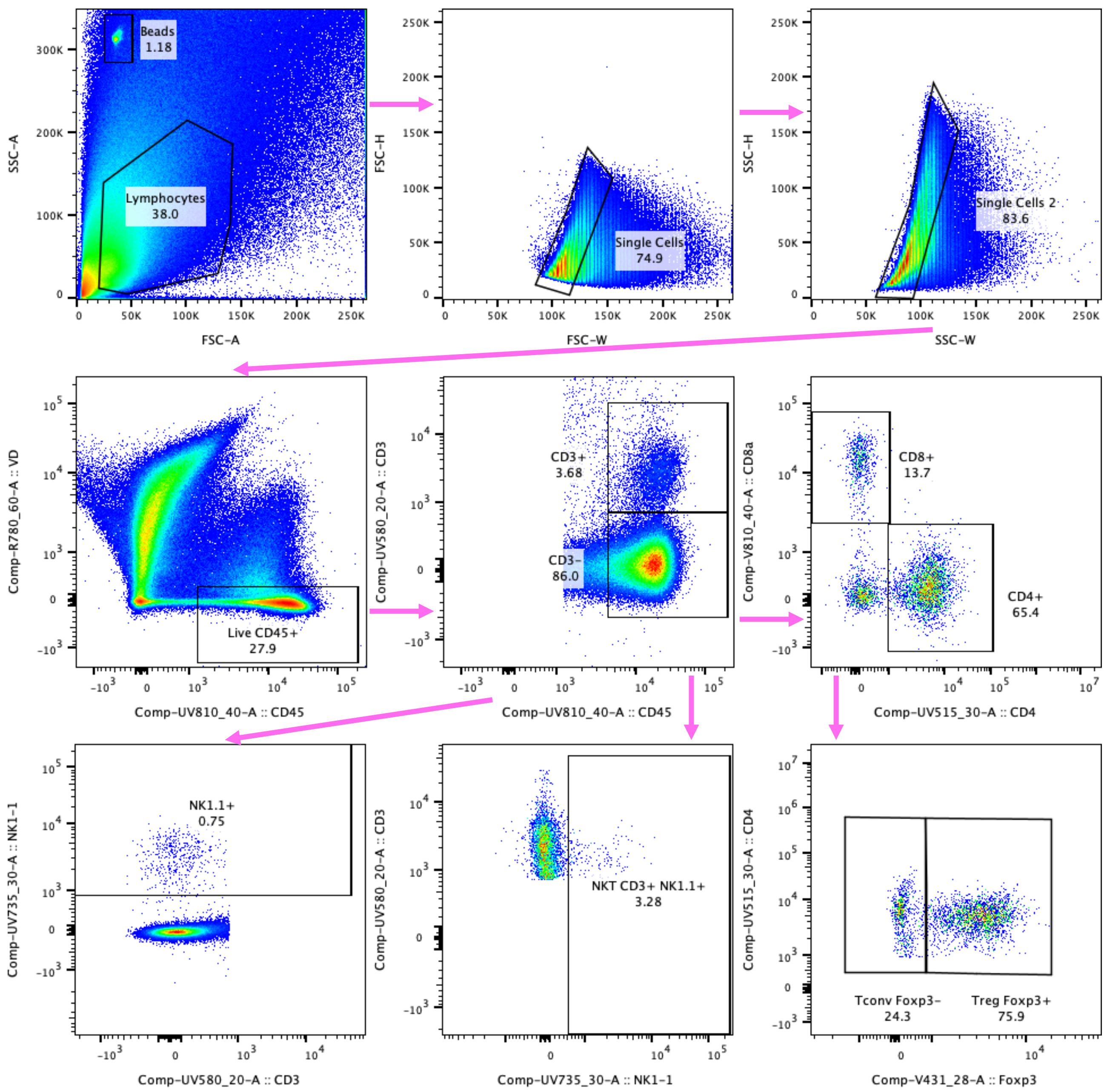

Beads
1.18
Lymphocytes
38.0
SSC-A
FSC-A
FSC-H
FSC-W
Single Cells
74.9
SSC-H
SSC-W
Single Cells 2
83.6
Comp-R780_60-A :: VD
Comp-UV810_40-A :: CD45
Live CD45+
27.9
Comp-UV580_20-A :: CD3
CD3+
3.68
CD3-
86.0
Comp-V810_40-A :: CD8a
Comp-UV515_30-A :: CD4
CD8+
13.7
CD4+
65.4
Comp-UV735_30-A :: NK1-1
Comp-UV580_20-A :: CD3
NK1.1+
0.75
Comp-UV735_30-A :: NK1-1
NKT CD3+ NK1.1+
3.28
Comp-UV515_30-A :: CD4
Comp-V431_28-A :: Foxp3
Tconv Foxp3-
24.3
Treg Foxp3+
75.9

**Fig. S11: Wild-type lymphocyte panel gating strategy.**

Live single cells were gated for $CD45^+$ expression, then classified as: Tconv, $TCR\beta^+$ $CD4^+$ $Foxp3^-$; Tregs, $TCR\beta^+$ $CD4^+$ $Foxp3^+$; and $CD8^+$ T-cells, $TCR\beta^+$ $CD8^+$.

### Figure S12

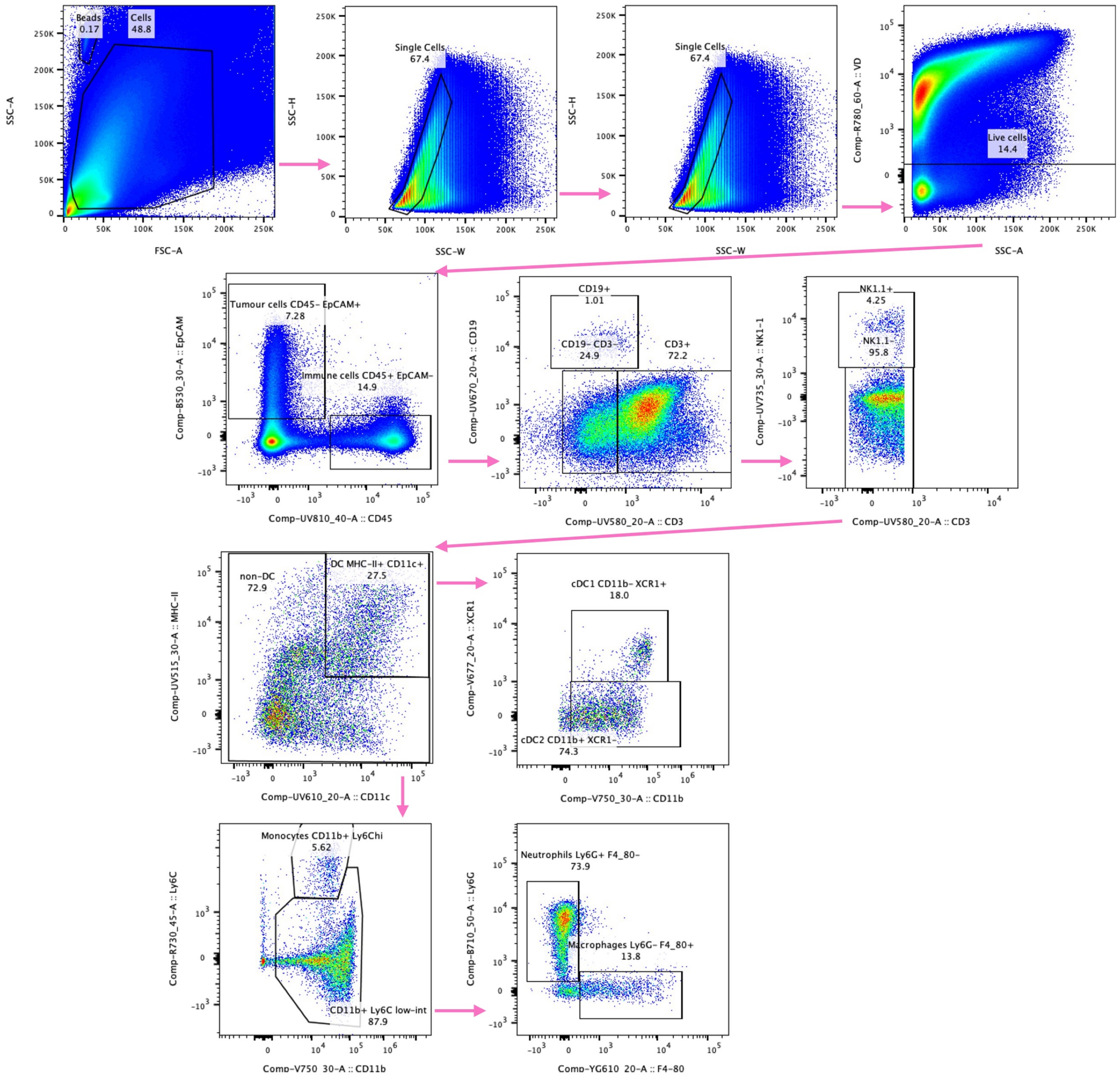
Beads
0.17
Cells
48.8
SSC-A
FSC-A
Single Cells
67.4
SSC-H
SSC-W
Single Cells
67.4
SSC-H
SSC-W
Live cells
14.4
Comp-R780_60-A :: VD
SSC-A
Tumour cells CD45- EpCAM+
7.28
Immune cells CD45+ EpCAM-
14.9
Comp-B530_30-A :: EpCAM
Comp-UV810_40-A :: CD45
CD19+
1.01
CD19- CD3-
24.9
CD3+
72.2
Comp-UV670_20-A :: CD19
Comp-UV580_20-A :: CD3
NK1.1+
4.25
NK1.1-
95.8
Comp-UV735_30-A :: NK1-1
Comp-UV580_20-A :: CD3
non-DC
72.9
DC MHC-II+ CD11c+
27.5
Comp-UV515_30-A :: MHC-II
Comp-UV610_20-A :: CD11c
cDC1 CD11b- XCR1+
18.0
cDC2 CD11b+ XCR1-
74.3
Comp-V677_20-A :: XCR1
Comp-V750_30-A :: CD11b
Monocytes CD11b+ Ly6Chi
5.62
CD11b+ Ly6C low-int
87.9
Comp-R730_45-A :: Ly6C
Comp-V750_30-A :: CD11b
Neutrophils Ly6G+ F4_80-
73.9
Macrophages Ly6G- F4_80+
13.8
Comp-B710_50-A :: Ly6G
Comp-YG610_20-A :: F4-80

**Fig. S12: Myeloid panel gating strategy.**

All gated from live single cells, $CD45^{+}$ $EpCAM^{-}$ $CD19^{-}$ $CD3^{-}$. NK cells: $CD3^{-}$, $NK1.1^{+}$.DCs: $NK1.1^{-}$, $MHC\text{-}II^{+}$ $CD11c^{+}$; cDC1: $MHC\text{-}II^{+}$ $CD11c^{+}$ $XCR1^{+}$; cDC2: $MHC\text{-}II^{+}$ $CD11c^{+}$ $XCR1^{-}$; Neutrophils: $CD11b^{+}$ $Ly6C^{low\text{-}int}$ $Ly6G^{high}$, Monocytes: $CD11b^{+}$ $Ly6C^{hi}$, Macrophages: $MHC\text{-}II^{+}$ $CD11b^{+}$ $Ly6C^{low\text{-}int}$ $F4/80^{+}$; M1 Macrophages: $MHC\text{-}II^{+}$ $CD11b^{+}$ $Ly6C^{low\text{-}int}$ $F4/80^{+}$ $CD206^{-}$, M2 Macrophages: $MHC\text{-}II^{+}$ $CD11b^{+}$ $Ly6C^{low\text{-}int}$ $F4/80^{+}$ $CD206^{+}$.

### Figure S13

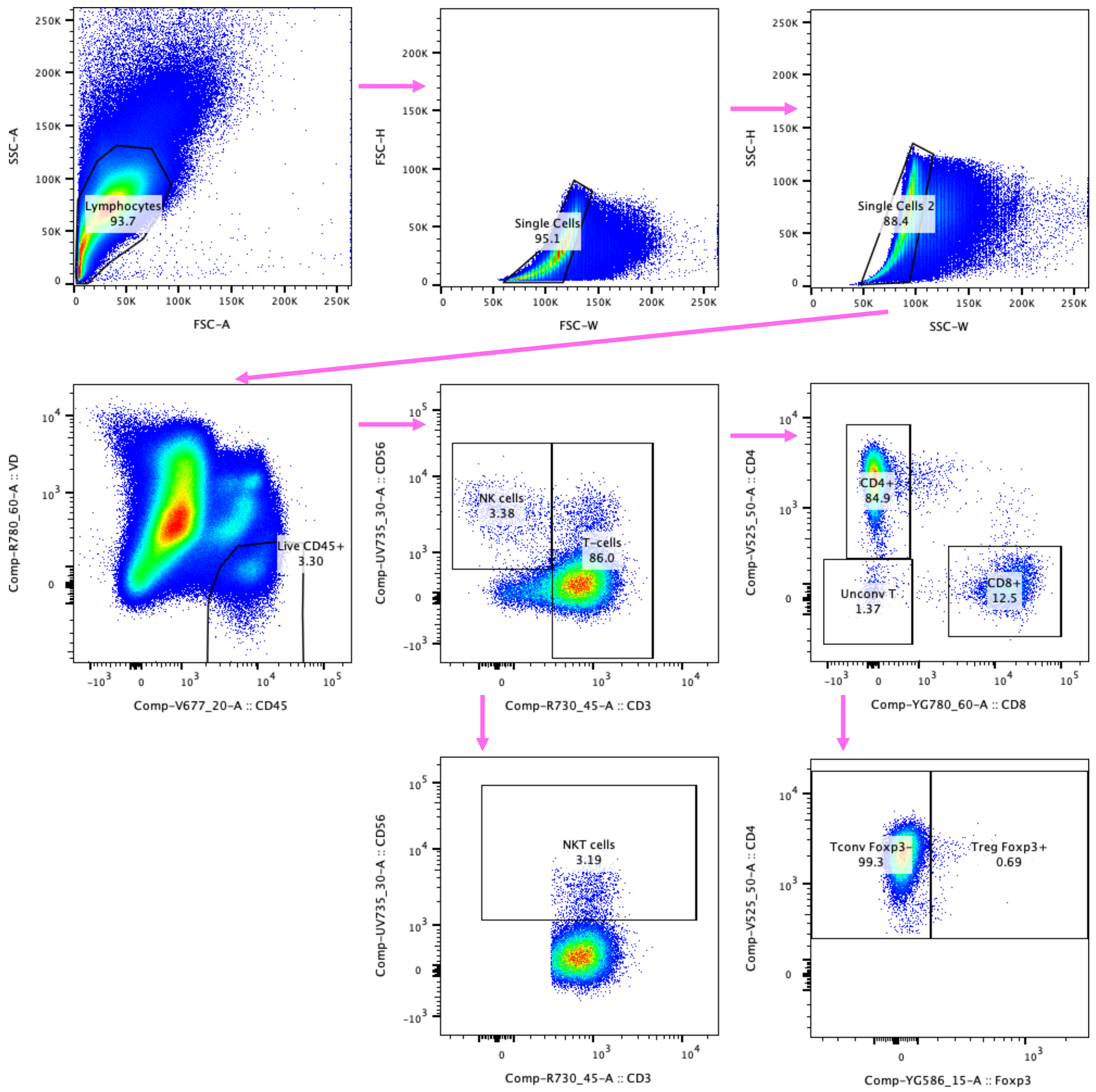

SSC-A
FSC-A
Lymphocytes
93.7
FSC-H
FSC-W
Single Cells
95.1
SSC-H
SSC-W
Single Cells 2
88.4
Comp-R780_60-A :: VD
Comp-V677_20-A :: CD45
Live CD45+
3.30
Comp-UV735_30-A :: CD56
Comp-R730_45-A :: CD3
NK cells
3.38
T-cells
86.0
Comp-V525_50-A :: CD4
Comp-YG780_60-A :: CD8
CD4+
84.9
Unconv T
1.37
CD8+
12.5
NKT cells
3.19
Tconv Foxp3-
99.3
Treg Foxp3+
0.69
Comp-YG586_15-A :: Foxp3

**Fig. S13: Human PBMC lymphocyte gating strategy.**

Live single cells were gated for $CD45^+$ expression, then classified as: NK cells, $CD3^-$ $CD56^+$; and T-cells, $CD3^+$ $CD56^-$ which were subclassified into: $CD4^+$ $Foxp3^-$; Tregs, $CD4^+$ $Foxp3^+$; and $CD8^+$ T-cells, $CD8^+$; NKT cells, $CD56^+$; and unconventional T-cells, $CD4^-$ $CD8^-$.

### Figure S14

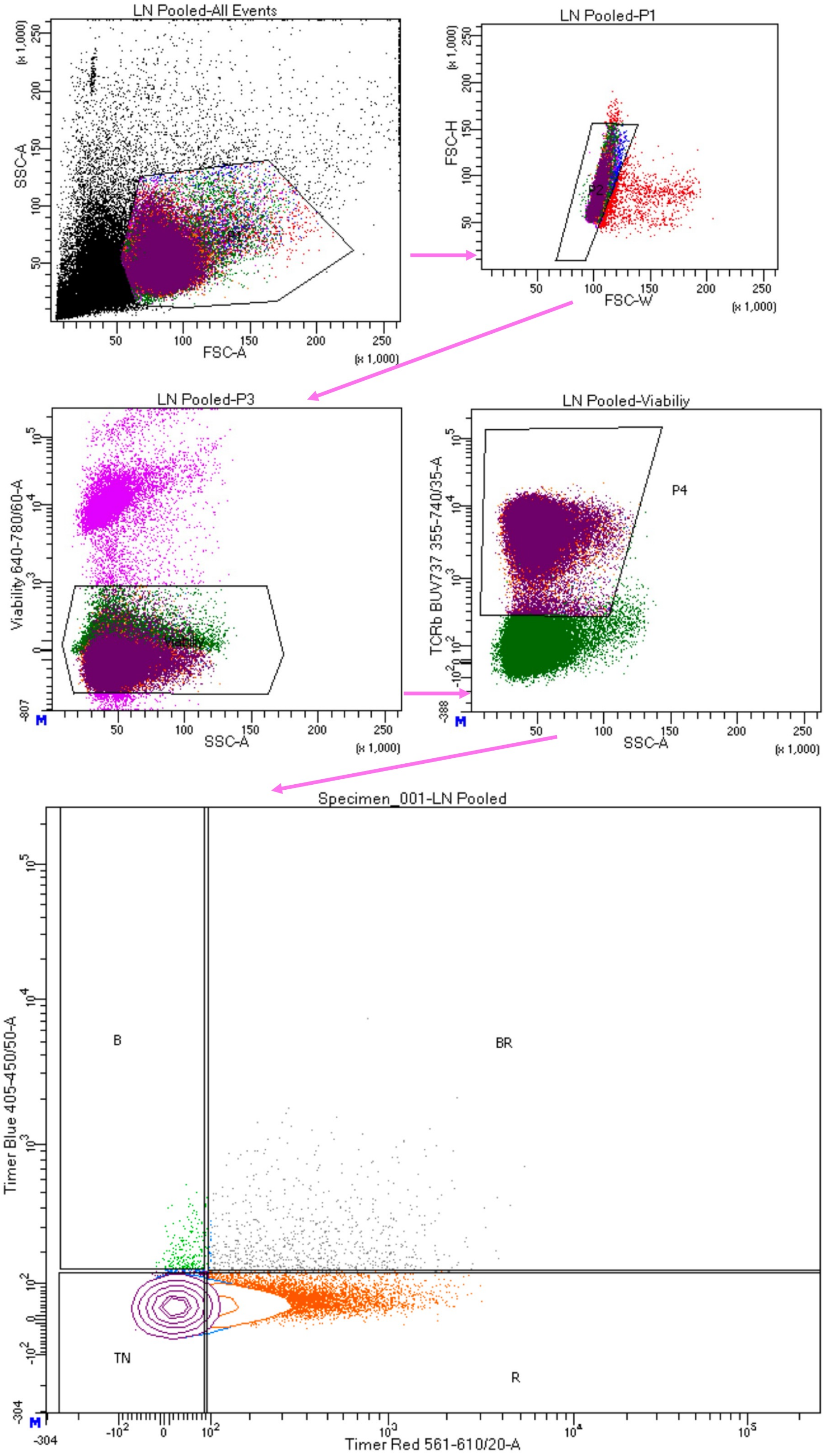


Tube: LN Pooled

| Population | #Events | %Parent | %Total |
|---|---|---|---|
| All Events | 215,502 | #### | 100.0 |
| P1 | 113,122 | 52.5 | 52.5 |
| P2 | 111,689 | 98.7 | 51.8 |
| P3 | 110,513 | 98.9 | 51.3 |
| Viabiliy | 100,000 | 90.5 | 46.4 |
| P4 | 43,105 | 43.1 | 20.0 |
| B | 243 | 0.6 | 0.1 |
| BR | 806 | 1.9 | 0.4 |
| R | 12,173 | 28.2 | 5.6 |
| TN | 29,823 | 69.2 | 13.8 |

**Fig. S14: Cell sorter gating strategy for single-cell sequencing.**

Live single cells were gated for TCRβ$^+$ expression and separated into four Tocky fluorescent timer (FT) protein populations: B (“new”), FT blue$^+$ FT red$^-$; BR (“persistent”), FT blue$^+$ FT red$^+$; R (“arrested”), FT blue$^-$ FT red$^+$; and TN (“timer negative”), FT blue$^-$ FT red$^-$.

**Reagents, chemicals and *in vivo* antibodies**

The following reagents/chemicals were used: ketamine (100 mg/kg, KETAVET), xylazine (16 mg/kg, ROMPUN), cisplatin (Sigma Aldrich, P4394), collagenase type 76 I-S (Sigma-Aldrich, C1639), Dispase II protease (Sigma-Aldrich, D4693), DNase I (Roche, 10104159001), ACK lysis buffer (Thermo Fisher Scientific, A1049201), anti-mouse CD16/CD32 Fc-receptor blocker (BD, 553142), anti-human Fc-receptor blocker (BD, 564220), CTL Anti-Aggregate Wash Supplement (Cellular Technology Limited, CTL-AA-005), Foxp3/Transcription Factor Staining Buffer Set (Thermo Fisher Scientific, 00-5523-00), UltraComp eBeads compensation beads (Thermo Fisher Scientific, 01-2222-41), CountBright Absolute Counting Beads (Thermo Fisher Scientific, C36950), mouse CD45 (TIL) microbeads (Milentyi Biotec, 130-052-301), Fixable Viability Dye eFluor 780 (Thermo Fisher Scientific, 65-0865-14), Calcein AM (Thermo Fisher Scientific, C1430), DRAQ7 (BD, 564904), Buffer RLT Plus (Qiagen, 1053393), β-mercaptoethanol (Gibco, 21985023) and RNeasy Plus Mini Kit (Qiagen, 74134) with QIAshredder (Qiagen, 79656). The following *in vivo* antibodies were used: CD8a depletion antibody (BioXcell, clone: 2.43, cat no: BE0061, lot number: 811522A2), rat IgG2b isotype control (BioXcell, clone: LTF-2, cat no: BE0090, lot number: 831023M), anti-PD-1 antibody (BioXCell, clone: RMP1-14, cat no: BE0146, lot number: 810421N1) and rat IgG2a isotype control (clone: 2A3, BioXCell, cat no: BE0089, lot number: 796721M2). The following kits were used for single cell sequencing: BD Single-Cell Multiplexing Kit (BD, 633793), BD Rhapsody cDNA kit (BD, 633773), BD Rhapsody WTA Amplification kit (BD, 633801) and Mouse TCR/BCR Amplification Kit (BD, 666282).

**Table S1. List of anti-mouse antibodies used for immunoprofiling**

| Epitope | Clone | Fluorophore | Manufact-urer | Catalog number | Lot number | Dilution |
|---|---|---|---|---|---|---|
| **CD103** | M290 | BUV805 | BD | 741948 | 3095536 | 1:100 |
| **CD11b** | M1/70 | BV750 | Biolegend | 101267 | B441192 | 1:200 |
| **CD11c** | N418 | BUV615 | BD | 751222 | 5083061 | 1:100 |
| **CD19** | ID3 | BUV395 | BD | 565965 | 3214735 | 1:200 |
| **CD25** | PC61 | AF700 | Biolegend | 102024 | B365418 | 1:100 |
| **CD3** | 17A2 | BUV563 | BD | 741319 | 5147291 | 1:100 |
| **CD4** | RM4-5 | APC | Biolegend | 100515 | B288206 | 1:200 |
| **CD4** | RM4-4 | BUV496 | BD | 741051 | 1197023 | 1:200 |
| **CD4** | RM4-5 | PerCP | Biolegend | 100538 | B368726 | 1:100 |
| **CD44** | IM7 | BUV395 | BD | 568507 | 4288992 | 1:100 |
| **CD45** | 30-F11 | AF700 | Biolegend | 103128 | B274307 | 1:300 |
| **CD45** | 30-F11 | BUV805 | BD | 568336 | 4354579 | 1:300 |
| **CD62L** | MEL-14 | BV510 | Biolegend | 104441 | B403539 | 1:100 |
| **CD69** | H1.2F3 | PE-Cy7 | Biolegend | 104512 | B330013 | 1:100 |
| **CD8α** | 53-6.7 | BV785 | Biolegend | 100750 | B384483 | 1:100 |
| **CD8α** | 53-6.7 | PE-Cy7 | Biolegend | 100721 | B282417 | 1:300 |
| **CD8β** | 53-5.8 | FITC | Biolegend | 140403 | B274128 | 1:200 |
| **CD80** | 16-10A1 | BV510 | Biolegend | 104741 | B349630 | 1:100 |
| **CD86** | GL-1 | APC | Biolegend | 105011 | B346110 | 1:100 |
| **EpCAM** | G8.8 | AF488 | Biolegend | 118210 | B337007 | 1:200 |
| **F4/80** | BM8 | PE/Dazzle | Biolegend | 123145 | B375240 | 1:100 |
| **FOXP3** | FJK-16s | eFluor 450 | eBioscience | 48-5773-82 | 2401080 | 1:100 |
| **Granzyme B** | GB11 | AF647 | Biolegend | 515405 | B266742 | 1:100 |
| **Ly6C** | HK1.4 | AF700 | Biolegend | 128023 | B388651 | 1:100 |
| **Ly6G** | 1A8 | PerCP/Cy5.5 | Biolegend | 127615 | B380284 | 1:100 |
| **MHC-II** | M5/114.15.2 | BUV496 | BD | 750281 | 4080119 | 1:200 |
| **NK1.1** | PK136 | BUV737 | BD | 741715 | 4047857 | 1:100 |
| **PD-1** | RMP1-30 | APC | Biolegend | 109112 | B384618 | 1:100 |
| **PD-1** | 29F.1A12 | BV605 | Biolegend | 135219 | B333822 | 1:200 |
| **Perforin** | S16009B | PE | Biolegend | 154405 | B402167 | 1:100 |
| **T-bet** | 4B10 | PE-Cy7 | eBioscience | 25-5825-82 | 2410093 | 1:100 |
| **TCRβ** | 29A1.4 | BUV737 | BD | 612821 | 1315739 | 1:300 |
| **TIM-3** | RMT3-23 | BV711 | Biolegend | 119727 | B362528 | 1:100 |
| **XCR1** | ZET | BV650 | Biolegend | 148220 | B430650 | 1:200 |

**Table S2. List of anti-human antibodies used for immunoprofiling**

| Epitope | Clone | Fluorophore | Manufact-urer | Catalog number | Lot number | Dilution |
|---|---|---|---|---|---|---|
| **CCR7** | G043H7 | BUV395 | eBioscience | 363-1979-41 | 2996996 | 1:17 |
| **CD25** | BC96 | RB744 | BD | 570482 | 3251820 | 1:17 |
| **CD27** | O323 | BUV805 | eBioscience | 368-0279-41 | 2976777 | 1:17 |
| **CD3** | HIT3a | AF700 | Biolegend | 300324 | B368722 | 1:100 |
| **CD4** | SK3 | BV510 | Biolegend | 344634 | B350564 | 1:200 |
| **CD45** | HI30 | BV650 | Biolegend | 304044 | B343579 | 1:200 |
| **CD45RA** | HI100 | FITC | Biolegend | 304105 | B314028 | 1:100 |
| **CD45RO** | UCHL1 | BV421 | Biolegend | 302630 | B304216 | 1:50 |
| **CD56** | TULY56 | BUV737 | BD | 569592 | 5078607 | 1:17 |
| **CD62L** | DREG-56 | PerCP Cy5.5 | Biolegend | 304824 | B268797 | 1:100 |
| **CD69** | FN50 | PE-Cy5 | Biolegend | 310907 | B385121 | 1:17 |
| **CD8** | SK1 | PE-Cy7 | Biolegend | 344712 | B375218 | 1:200 |
| **Foxp3** | PCH101 | PE | eBioscience | 12-4776-41 | 2970942 | 1:40 |
| **Granzyme B** | GB11 | PE-CF594 | BD | 562462 | 4326373 | 1:100 |
| **Ki67** | B56 | RB670 | BD | 571924 | 4261363 | 1:17 |
| **PD-1** | EH12.2H7 | APC | Biolegend | 329908 | B362224 | 1:17 |
| **Perforin** | dG9 | BV711 | Biolegend | 308129 | B428509 | 1:17 |
| **TCF7/TCF1** | S33-966 | RB613 | BD | 571352 | 5091273 | 1:17 |
| **TIM-3** | F38-2E2 | BV785 | Biolegend | 345032 | B350351 | 1:17 |
| **TOX** | NAN448B | RB780 | BD | 570193 | 5008032 | 1:17 |

**Table S3. BD Abseq antibodies**

| | BD AbSeq Antibody-Oligo target | Clone | Catalog number | Lot number |
|---|---|---|---|---|
| 1 | CD103 | 2E7 | 940358 | 3045960 |
| 2 | CD137 (4-1BB) | 1AH2 | 940199 | 2305124 |
| 3 | CD154 (CD40L) | MR1 | 940435 | 2132437 |
| 4 | CD223 (LAG3) | C9B7W | 940152 | 2110971 |
| 5 | CD224.2 (2B4 B6 alloantigen) | 2B4 | 940474 | 3066364 |
| 6 | CD25 (IL-2 Receptor α) | PC61 | 940116 | 2132304 |
| 7 | CD27 | LG.3A10 | 940157 | 2214562 |
| 8 | CD278 (ICOS) | 7E.17G9 | 940176 | 3045981 |
| 9 | CD279 (PD-1) | RMP1-30 | 940343 | 2132266 |
| 10 | CD28 | 37.51 | 940120 | 3026297 |
| 11 | CD366 (TIM-3) | 5D12 | 940207 | 3026279 |
| 12 | CD38 | 90/CD38 | 940162 | 3059103 |
| 13 | CD4 | RM4-5 | 940108 | 2257213 |
| 14 | CD44 | IM7 | 940114 | 3066370 |
| 15 | CD62L (L-selectin) | MEL-14 | 940122 | 3012350 |
| 16 | CD69 | H1.2F3 | 940126 | 2222349 |
| 17 | CD8a | 53-6.7 | 940345 | 2074465 |
| 18 | KLRG1 | 2F1 | 940146 | 2305123 |
| 19 | NKG2A/C/E | 20D5 | 940444 | 3074560 |
| 20 | TIGIT | 1G9 | 940191 | 2293547 |
